# Determinants of FtsZ C-terminal linker-dependent regulation of cell wall metabolism in *Caulobacter crescentus*

**DOI:** 10.1101/632075

**Authors:** Kousik Sundararajan, Jordan M. Barrows, Erin D. Goley

**Affiliations:** Department of Biological Chemistry, Johns Hopkins University School of Medicine, Baltimore, MD 21205; Department of Biochemistry, Stanford University School of Medicine, Stanford, CA 94305

**Keywords:** *Caulobacter*, bacterial cell division, Z-ring, FtsZ, intrinsic disorder, divisome

## Abstract

Bacterial cell division requires assembly of a multi-protein machinery or “divisome” that remodels the cell envelope to cause constriction. The cytoskeletal protein FtsZ forms a ring-like scaffold for the divisome at the incipient division site. FtsZ has three major regions – a conserved, polymerizing GTPase domain; a C-terminal conserved (CTC) peptide required for binding membrane-anchoring proteins; and a C-terminal linker (CTL) of poor length and sequence conservation. We previously demonstrated that, in *Caulobacter crescentus*, the CTL regulates FtsZ polymerization *in vitro* and cell wall metabolism *in vivo*. To understand the mechanism of CTL-dependent regulation of cell wall metabolism, here we investigated the impact of the CTL on Z-ring structure in cells and employed genetics to identify molecular determinants of the dominant lethal effects of ΔCTL. Deleting the CTL specifically resulted in formation of dense, asymmetric, non-ring FtsZ assemblies *in vivo*. Moreover, we observed that production of an FtsZ variant with the GTPase domain of *Escherichia coli* FtsZ fused to the CTC of *C. crescentus* FtsZ phenocopied the effects of *C. crescentus* ΔCTL, suggesting the CTC mediates signaling to cell wall metabolism. Finally, whereas overproduction of ZapA, FzlC, or FtsEX had slight protective effects against ΔCTL, depletion of FtsA partially suppressed the effects of ΔCTL. From these results, we propose that the cell wall misregulation downstream of ΔCTL results from its aberrant assembly properties and is propagated through the interaction between the CTC of FtsZ and FtsA. Our study provides mechanistic insights into CTL-dependent regulation of cell wall enzymes downstream of FtsZ.

**Importance:** Bacterial cell division is essential and requires the recruitment and regulation of a complex network of proteins needed to initiate and guide constriction and cytokinesis. FtsZ serves as a master regulator for this process, and its function is highly dependent on both its self-assembly into a canonical “Z-ring” and interaction with protein binding partners, which results in the activation of enzymes that remodel the cell wall to drive constriction. Using mutants of FtsZ and its binding partners, we have established the role of its C-terminal linker domain in regulating Z-ring organization, as well as the requirement for its C-terminal conserved peptide and interaction with the membrane-anchoring protein FtsA for regulating cell wall remodeling for constriction.

## Introduction

Bacterial cell division requires spatially- and temporally-coordinated remodeling of the cell envelope to cause constriction. To this end, a multi-protein machinery called the divisome is assembled at the incipient site of division. The first and most conserved protein of the divisome, FtsZ, is a tubulin homolog that polymerizes into a discontinuous ring-like scaffold or “Z-ring” for the recruitment of other members of the divisome (1). Over two-dozen proteins are directly or indirectly recruited to the divisome in an FtsZ-dependent manner. In *Caulobacter crescentus*, these include FtsZ-binding proteins that regulate Z-ring structure (ZapA-ZauP, FzlA), membrane-anchoring proteins of FtsZ (FtsA, FzlC, FtsEX), cell wall enzymes (DipM, AmiC, FtsI/Pbp3, FtsW) and their regulators (FtsN, FtsQLB), outer membrane remodeling proteins (Tol-Pal complex), polarity factors (TipN), and proteins involved in chromosome segregation and translocation (FtsK) (2–3). While most of the essential members of the divisome have likely been identified, the interactions among these proteins and the regulation of their organization and function are unclear.

In addition to serving as a scaffold, FtsZ regulates the dynamic movement and activity of cell wall enzymes in the divisome, at least in some bacteria. Recent studies have shown that clusters of FtsZ protofilaments in the Z-ring undergo treadmilling motion that drives the movement of cell wall enzymes in *Escherichia coli* and *Bacillus subtilis* (5–6). This FtsZ-dependent regulation of cell wall enzyme dynamics is important for septum morphology in *E. coli* and for defining the rate of cell wall synthesis in *B. subtilis.* Moreover, in *Caulobacter crescentus*, we engineered FtsZ mutants that can assemble at midcell, recruit the divisome, and drive local cell wall synthesis, but that nevertheless cause lethal defects in cell wall metabolism and division failure (4). These observations suggest that cell division requires FtsZ-dependent regulation of the activity of the divisome-associated cell wall enzymes. Collectively these data indicate that Z-ring assembly properties are directly relevant to the regulation of local cell wall remodeling. However, the pathways downstream of Z-ring assembly that regulate cell wall enzymes are largely unknown.

FtsZ has three regions: (i) a conserved GTPase domain, (ii) a C-terminal linker (CTL), and (iii) a conserved C-terminal peptide (CTC) (Figure S1) (7). The GTPase domain is structurally similar to eukaryotic tubulin (8–9) and is sufficient for polymerization on binding GTP (4, 10). Mutations in the GTPase domain affect Z-ring dynamics, organization, and regulation of cell wall synthetic enzymes, at least in some bacteria (5–6). The CTC is composed of a conserved α-helix that is required for interaction of FtsZ with membrane-anchoring proteins such as FtsA (across multiple species of bacteria) and FzlC (in *C. crescentus*). The CTL is an intrinsically disordered region that connects the GTPase domain to the CTC and varies in length and sequence across species. While there are no known binding partners for the CTL, changes in length and sequence of the CTL affect polymer turnover and lateral interactions between FtsZ protofilaments, at least in the cases of FtsZ from *Caulobacter crescentus*, *Bacillus subtilis*, and *Agrobacterium tumefaciens* (10–12). Surprisingly, large modifications of CTL sequence are tolerated in *B. subtilis* and *E. coli* cells as long as flexibility of the CTL and a length range ± 50% of WT length are maintained (11, 13). Conversely, in *C. crescentus*, large truncations of the CTL are tolerated to some extent, but significant changes to CTL sequence impact protein stability and, therefore, cell division (4). Complete deletion of the CTL causes dominant lethal defects in Z-ring assembly and cell lysis, at least in *C. crescentus* and *B. subtilis* (4, 11). Identifying the contributions of the CTL to FtsZ function is essential to understanding the communication between Z-ring structure and cell wall enzyme activities.

We previously showed that the expression of FtsZ lacking its CTL (“ΔCTL”, wherein the GTPase domain is fused directly to the CTC) in the α-proteobacterium *C. crescentus* causes misregulation of cell wall enzymes resulting in the formation of spherical envelope bulges at the sites of ΔCTL assembly and rapid cell lysis (4). Using a fluorescent fusion to ZapA, a protein that binds FtsZ, we found that FtsZ superstructure was affected in ΔCTL: ΔCTL formed large, amorphous assemblies instead of focused rings (4). FtsZ with a minimal CTL of 14 amino acids (L14) exhibited WT-like Z-ring shape and did not lead to bulging and lysis. *In vitro*, ΔCTL polymerizes into straight multi-filament bundles that are significantly longer than the curved protofilaments observed for WT FtsZ or L14 by electron microscopy (4, 10). Moreover, ΔCTL exhibits lower GTP hydrolysis rates, reduced polymer turnover, and increased protofilament lateral interactions compared to WT FtsZ *in vitro* (4, 10, 14). These effects result in the formation of stable networks of ΔCTL protofilaments on membranes, in contrast to small dynamic clusters formed by WT FtsZ, when observed on supported lipid bilayers by total internal reflection fluorescence microscopy (14). Unlike the CTL, the CTC does not significantly contribute to polymer structure or dynamics for *C. crescentus* FtsZ – polymer structure, observed by EM, or GTP hydrolysis rates of FtsZ lacking its CTC (ΔCTC) are comparable to those of WT FtsZ (10). *In vivo*, ΔCTC forms Z-rings similar to WT but is incapable of cytokinesis (4).

Determining how CTL-dependent changes in FtsZ polymerization are communicated to the divisome is essential for understanding how Z-ring structure and dynamics regulate cell wall metabolism. We hypothesize that there are specific pathways downstream of FtsZ that contribute to the misregulation of cell wall enzymes caused by aberrant ΔCTL superstructures observed *in vitro* and *in vivo*. In the current study, we tested the effects of candidate division proteins on the lethal cell wall metabolic defects downstream of ΔCTL. Of all the division proteins tested, only FtsA appears to be required for ΔCTL-induced bulging and lysis. Specifically, we observed that a temperature sensitive allele of *ftsA* eliminated the bulging induced by ΔCTL. By expressing chimeric FtsZs bearing domains from *E. coli* and/or *C. crescentus* FtsZ in *C. crescentus* cells, we found that a chimeric FtsZ with the *E. coli* GTPase domain and *C. crescentus* CTC causes bulging and lysis but only in the absence of a CTL from either organism. Together, our results suggest that the CTL is required for proper Z-ring assembly, and the interaction between the CTC and FtsA is required for CTL-dependent signaling from Z-ring structure and/or dynamics to the regulation of cell wall enzymes in cells.

## Results and Discussion

### The CTL of FtsZ impacts Z-ring superstructure *in vivo*

We previously observed that the CTL affects the higher order assembly of FtsZ polymers *in vitro* (10, 14). Z-ring structure also appears to be regulated in a CTL-dependent manner in cells when imaged using a fluorescently-labeled FtsZ binding protein (ZapA-Venus) (4). To test if the differences in ZapA-Venus structures observed previously reliably reflect differences in Z-ring organization, we directly visualized N-terminal monomeric-NeonGreen (mNG) fluorescent fusions to FtsZ or ΔCTL using epifluorescence microscopy. We found that mNG-ΔCTL produced from a xylose-inducible promoter (P_*xylX*_) in the presence or absence of WT FtsZ caused filamentation, local envelope bulges, and rapid cell lysis, indicating that mNG-ΔCTL is dominant lethal, similar to untagged ΔCTL (Figures 1A, S2, S5). This allowed us to compare structures formed by FtsZ or ΔCTL *in vivo* to identify CTL-dependent differences in the Z-ring that might correlate with bulging and lysis. In our analysis, we also included mNG fusions to (i) L14 – an FtsZ CTL variant with a truncated 14 amino acid CTL that is incapable of cytokinesis but does not cause bulging and lysis, and (ii) *Hn*CTL – an FtsZ variant with the *Cc*CTL sequence replaced with the CTL from *Hyphomonas neptunium* FtsZ that causes inefficient cytokinesis (elongation and slower doubling time), as controls (4) (Figure S1).

**Figure 1:**
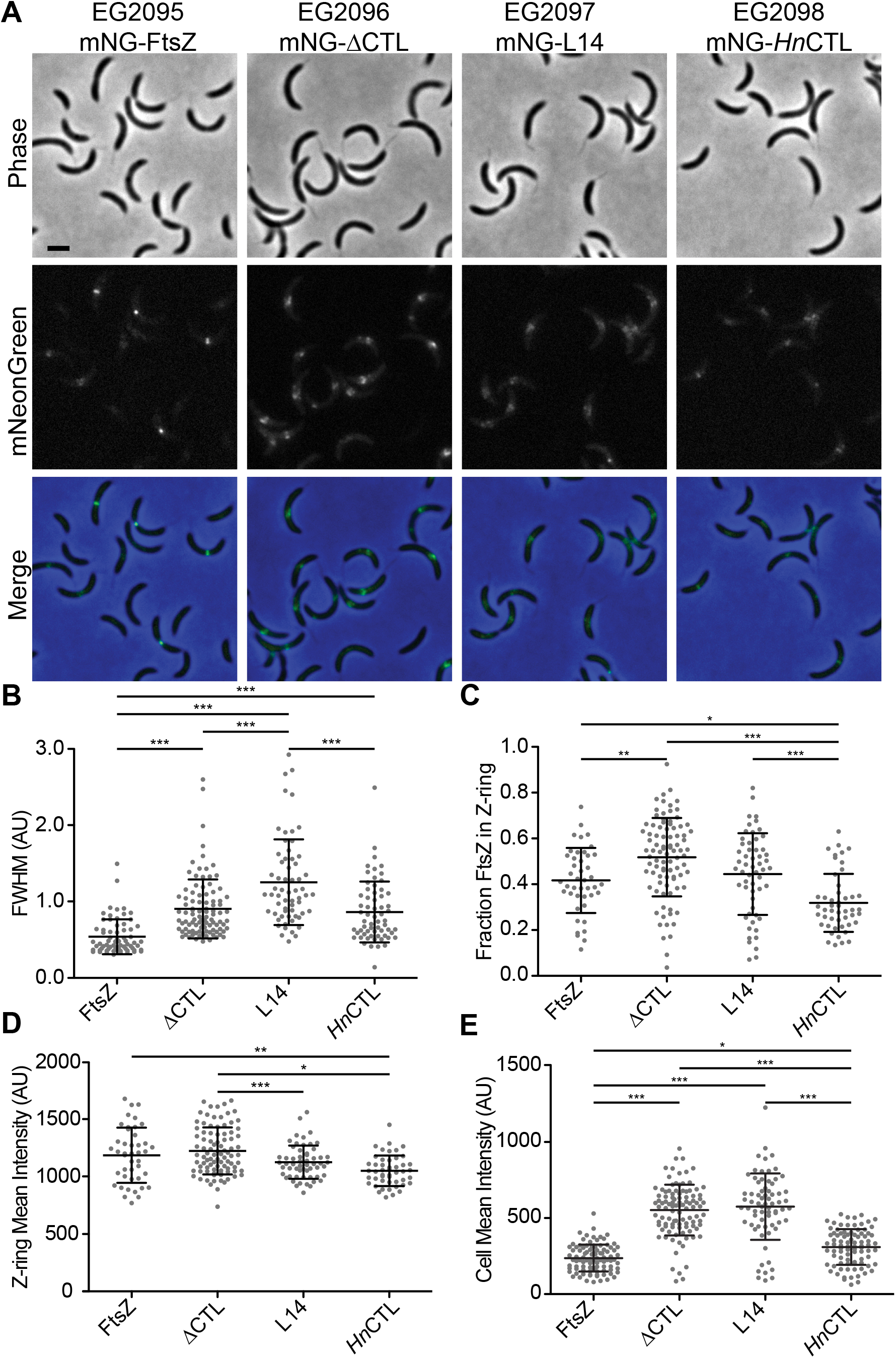
ΔCTL assembles into large asymmetric superstructures at sites of cell wall bulging in cells depleted of WT FtsZ. **A.** Phase contrast, epifluorescence, and merged images of cells induced with xylose to drive expression of *mNG-FtsZ, mNG-ΔCTL, mNG-L14*, or *mNG-*Hn*CTL* from the P_*xylX*_promoter for 1 hour while simultaneously depleting WT FtsZ. Scale bar – 2 µm. **B.-E.** Quantification of epifluorescence images of cells 3 to 5 µm long indicating the full-width at half max (FWHM) values of Z-ring intensity (**B)**, fraction of mNG-FtsZ or variants in the Z-ring (**C**), and mean epifluorescence intensity of the Z-ring (**D**) or the entire cells (**E**) in an FtsZ depletion background. Bars represent standard deviation. * - P ≤ 0.05; ** - P ≤ 0.01; *** - P ≤ 0.001. Strain key: mNG-FtsZ (EG2095), mNG-ΔCTL (EG2096), mNG-L14 (EG2097), mNG-*Hn*CTL (EG2098)

We expressed mNG fusions to FtsZ or CTL variants using the xylose-inducible P_*xylX*_promoter while simultaneously depleting WT FtsZ using strains wherein the only copy of *ftsZ* is under the control of the vanillate-inducible P_*vanA*_promoter. mNG-FtsZ formed ring-like structures that appear as a band or two closely spaced foci aligned along the short axis or a single focus per dividing cell after 1 hour of induction (Figure 1A). At longer induction times, mNG-FtsZ Z-ring structure was maintained despite cell filamentation due to depletion of WT FtsZ (Figure S2).

Unlike WT FtsZ, within 1 hour of induction, mNG-ΔCTL formed one or more wider and less ring-like structures (Figure 1A). These structures increased in size and intensity over time (Figure S2). Frequently, mNG-ΔCTL structures appeared to be asymmetrically distributed along the short axis of the cell (Figure 1A, S2). mNG-L14 assembled into apparently less dense, diffuse structures after 1 hour of induction (Figure 1A) which became more diffuse and scattered at longer induction times (Figure S2). mNG-*Hn*CTL structures appeared predominantly as faint rings or foci or more dispersed structures similar to mNG-L14, and did not change significantly with longer induction or cell filamentation (Figure 1A, S2).

We quantitatively analyzed Z-ring intensities and structures using MicrobeJ (15) and Oufti (16). To avoid potential effects of cell length on Z-ring organization, we focused on cells 3-5 µm long, which we determined from demographs (Figure S3) to have stable Z-rings after induction of mNG-FtsZ or CTL variants for 1 hour. We calculated the full-width at half maximum (FWHM) value for mNG intensity along the longitudinal axis for each variant, as a measure of degree of focusing of the Z-ring. Z-rings formed by mNG-ΔCTL, mNG-L14, and mNG-*Hn*CTL were wider compared to those formed by mNG-FtsZ (Figure 1B). We asked if differences in the fraction of FtsZ present in the Z-ring might contribute to the altered Z-ring structures formed by different CTL variants by determining relative enrichment of fluorescence signal at the Z-ring compared to the rest of the cell. Indeed, cells expressing mNG-ΔCTL had a significantly greater proportion of fluorescence signal in the Z-ring than those expressing mNG-FtsZ (Figure 1C). The fraction of mNG-*Hn*CTL was lower than each of the other CTL variants, suggesting a lower tendency to assemble into polymers at the Z-ring, while that of mNG-L14 was similar to WT FtsZ (Figure 1C). We next measured the mean fluorescence intensity (i.e. density) of Z-rings in each strain to determine whether variant Z-rings were more or less diffuse than those formed by WT FtsZ. While mNG-FtsZ and mNG-ΔCTL had similar mean intensity, mNG-L14 and mNG*-Hn*CTL each formed less intense structures (Figure 1D), consistent with their apparent “dimness” in the images and our biochemical studies indicating that L14 and *Hn*CTL do not polymerize as robustly as FtsZ or CTL (4, 10).

To address whether protein levels are affected by CTL composition, we determined the relative amount of each tagged FtsZ variant per cell in each strain. Mean fluorescence intensity values for the whole cell were significantly increased in cells expressing mNG-ΔCTL or mNG-L14 compared to those expressing mNG-FtsZ or mNG-*Hn*CTL (Figure 1E), suggesting an increased concentration of protein for each of these variants. Using quantitative immunoblotting, we found that indeed, mNG-ΔCTL and mNG-L14 levels were ~5-fold higher than mNG-FtsZ or mNG-*Hn*CTL and that these levels increased relative to mNG-FtsZ over time (Figure S4). mNG-L14 was present at higher levels than mNG-ΔCTL, whereas mNG-*Hn*CTL had levels nearly equivalent to mNG-FtsZ (Figure S4). Since all mNG fusions were expressed using identical induction conditions, increased steady state protein levels are likely due to differences in post-translational stability.

Finally, we tested if the structures formed by the CTL variants were influenced by the presence of WT FtsZ by expressing mNG fusions in an otherwise WT strain i.e. without depleting WT FtsZ. All four variants formed ring-like structures at 1 hour of induction, with mNG-ΔCTL forming slightly wider and brighter rings (Figure S5A). However, after 5 hours of induction, mNG-ΔCTL structures became less ring-like and more asymmetric, while the structures formed by the other CTL variants appeared largely similar to mNG-FtsZ (Figure S5B). This observation is in accordance with the ability of ΔCTL to cause bulging and lysis earlier in the absence of WT FtsZ than in its presence (4) and with the propensity of ΔCTL and WT to form long, bundled copolymers *in vitro* (10). In all cells expressing ΔCTL, the appearance of aberrant structures preceded the appearance of cell envelope bulges. This indicates that aberrant Z-ring morphology in the absence of CTL is not a result of altered cell geometry, but rather inherent to the assembly properties of ΔCTL. Quantitation of Z-ring structures formed by and protein levels of each CTL variant in the presence of WT FtsZ after 1 hour of induction showed similar trends as during depletion of FtsZ, with the exception that the mean Z-ring intensity for mNG-L14 was similar to that of mNG-ΔCTL in the presence of WT FtsZ (Figure S5, S6).

We previously observed that, *in vitro, Hn*CTL and L14 form relatively few, unbundled filaments while ΔCTL forms bundled, more stable filaments when compared to WT FtsZ (4, 10). We therefore postulate that the dispersed, less dense structures formed by mNG-L14 and mNG-*Hn*CTL result from their reduced polymerization propensity while the aberrant, non-canonical structures formed by ΔCTL are a consequence of its increased tendency to form bundles. The increased fraction of ΔCTL in the Z-ring likely reflects the hyperstability of ΔCTL polymers, as observed *in vitro* (10, 14). The absence of asymmetric, non-ring structures for mNG-FtsZ or mNG-*Hn*CTL, even in filamentous cells, suggests that the CTL-dependent effects we observed on Z-ring structure are not due to cell length or morphology, but are specific to the assembly properties of each FtsZ variant. In addition, the finding that total mNG-ΔCTL and -L14 protein levels are increased compared to mNG-FtsZ implies that ΔCTL and L14 variants exhibit increased protein stability. However, the differences in Z-ring structures formed by mNG-ΔCTL and mNG-L14 indicates that increased protein concentration is not sufficient to explain the altered Z-ring morphology we observed in cells. The increased fraction of mNG-ΔCTL in the Z-ring and increased mean Z-ring intensity in the absence of WT FtsZ and its capacity to form persistently aberrant structures in the presence of WT distinguish ΔCTL from the L14 variant, implicating these characteristics in the downstream misregulation of cell wall metabolism specific to ΔCTL.

### *E. coli* GTPase domain is sufficient to cause bulging and lysis when fused to *C. crescentus* CTC

Next, we sought to determine the contributions of each region of FtsZ to the bulging and lysis phenotype by making chimeric FtsZ variants using the GTPase domain, CTL, and/or CTC regions of *E. coli* and/or *C. crescentus* FtsZ (Figure 2). We reasoned that, due to the low sequence homology and distinct binding partners of FtsZs from these organisms, *C. crescentus* FtsZ binding partners would not be able to bind and regulate *E. coli* FtsZ. We expressed chimeric FtsZ mutants in *C. crescentus* cells depleted of WT FtsZ (using the strain wherein the only copy of *ftsZ* is under vanillate-driven expression) using the xylose-inducible P_*xylX*_ promoter and followed their effects on cell morphology. Additionally, we imaged the incorporation of fluorescently labeled D-alanine (HADA) (17) to visualize regions of active cell wall metabolism.

**Figure 2:**
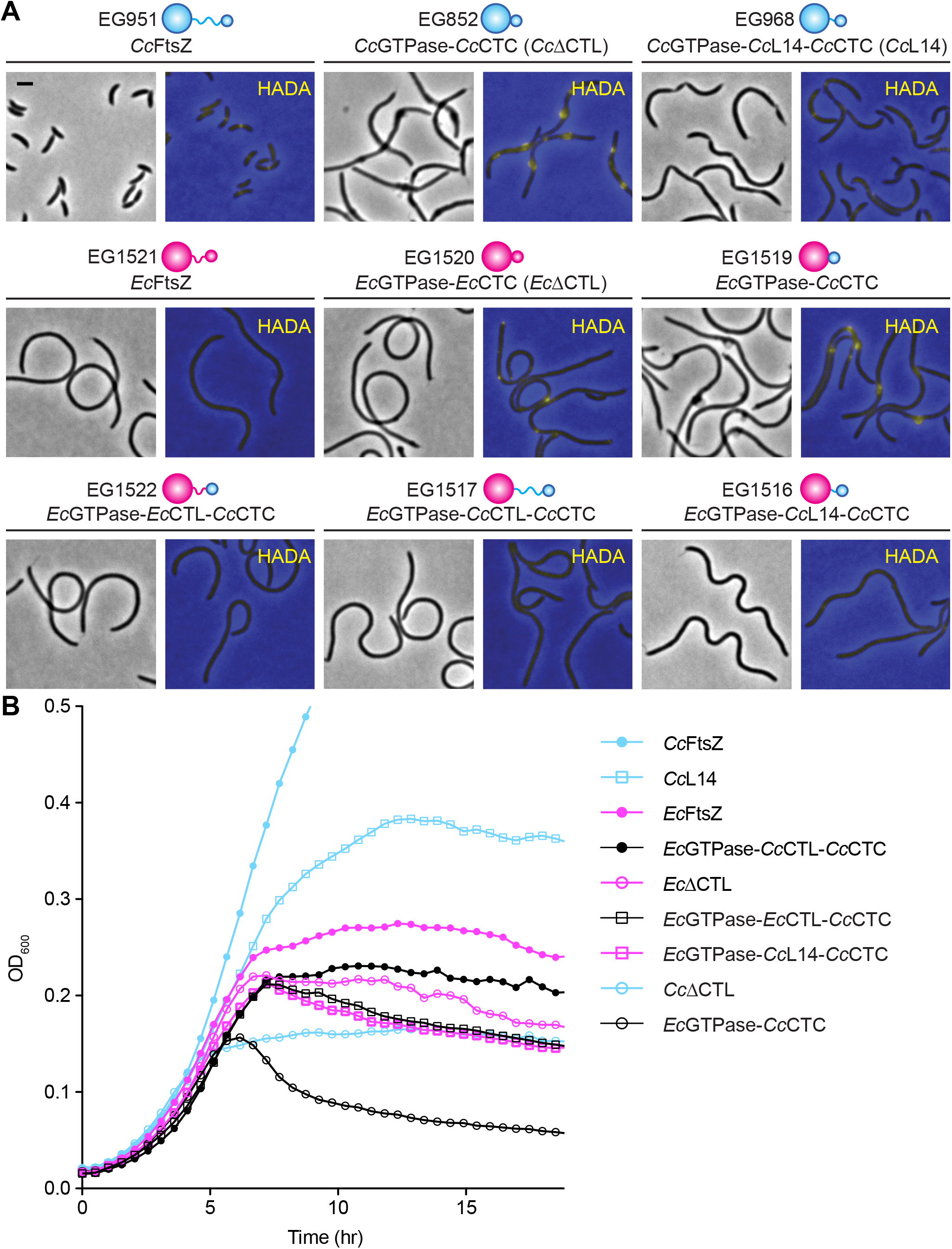
*Ec*GTPase-*Cc*CTC can cause bulging and lysis similar to *Cc*ΔCTL. **A.** Phase contrast images of morphology and merged epifluorescence images showing HADA incorporation (yellow) overlaid on phase contrast images (blue) corresponding to cells depleted of FtsZ and simultaneously induced for xylose-dependent production of *C. crescentus* FtsZ (cyan), *E. coli* FtsZ (magenta), CTL truncations, or their chimeric variants. Phase contrast images were acquired after 5 hours of induction of FtsZ variant. HADA fluorescence images were acquired after 4.5 hours of induction of FtsZ variant. Scale bar – 2 µm. **B.** Growth characteristics of cells in A. represented as absorbance at OD_600_ over time. Strain key: *Cc*FtsZ (EG951), *Cc*ΔCTL (EG852), *Cc*L14 (EG968), *Ec*FtsZ (EG1521), *Ec*ΔCTL (EG1520), *Ec*GTPase-*Cc*CTC (EG1519), *Ec*GTPase-*Ec*CTL-*Cc*CTC (EG1522), *Ec*GTPase-*Cc*CTL-*Cc*CTC (EG1517), *Ec*GTPase-*Cc*L14-*Cc*CTC (EG1516)

In *C. crescentus*, FtsZ drives the majority of cell wall synthesis at mid-cell (Figure 2A) and depletion of WT FtsZ in *C. crescentus* cells causes diffuse cell wall synthesis (18). As expected, xylose-induced production of *Cc*ΔCTL caused bulging and the bulges were sites of active cell wall synthesis (Figure 2A). On the other hand, xylose-induced *Cc*L14 could direct initiation of constriction and drive cell wall synthesis at multiple sites along filamentous cells, similar to the localization pattern of mNG-L14. Production of *Ec*FtsZ in *C. crescentus* cells did not result in any constriction or localized cell wall synthesis, consistent with the expectation that *Ec*FtsZ cannot efficiently engage the *C. crescentus* division or PG metabolic machinery. When we expressed EcΔ*CTL* (Ec*GTPase*-Ec*CTC*), we did not observe any constriction initiation, bulging, or lysis. However, this mutant was surprisingly able to drive limited local cell wall synthesis. The localization of HADA fluorescence appeared diffuse with occasional asymmetrically distributed foci along the short axis of the cells or at the cell pole.

Strikingly, when we expressed a chimera wherein *Ec*GTPase domain is fused to *Cc*CTC (*Ec*GTPase-*Cc*CTC), we observed cell envelope bulges similar to the effects of *Cc*ΔCTL. Once again, similar to *Cc*ΔCTL, bulges were the sites of active cell wall synthesis in these cells. Moreover, similar to the expression of CcΔ*CTL*, expression of Ec*GTPase*-Cc*CTC* resulted in rapid cell lysis (Figure 2B). The toxic effects of *Ec*GTPase-*Cc*CTC were not observed when we introduced either *Cc*CTL or *Ec*CTL back into this chimera, i.e., xylose-induced expression of either Ec*GTPase*-Cc*CTL*-Cc*CTC* or Ec*GTPase*-EcC*TL-CcCTC* resulted in smooth filamentous cells with diffuse cell wall synthesis, similar to cells with *Ec*FtsZ. Similar to introducing *Cc*CTL or *Ec*CTL, introducing *Cc*L14 (14 amino acids from the C-terminus of the *Cc*CTL) between *Ec*GTPase and *Cc*CTC (*Ec*GTPase-*Cc*L14-*Cc*CTC) did not cause constriction or cell envelope defects. We confirmed by immunoblotting using antibodies against both *C. crescentus* and *E. coli* FtsZ that there were no significant differences in the expression levels of these chimeras that could account for the differences in phenotypes observed (Figure S7).

Taken together, our results suggest that the peptidoglycan misregulation downstream of *Cc*ΔCTL assembly requires mainly three factors – (i) a polymerizing GTPase domain (4), (ii) absence of a minimal CTL, and (iii) *Cc*CTC. Moreover, since *Cc*ΔCTL (*Cc*GTPase-*Cc*CTC) and *Ec*GTPase-*Cc*CTC cause almost identical effects on cell morphology and cell wall integrity, and the *Cc*GTPase domain alone was previously shown to be insufficient to cause bulges (4), we conclude that interactions of FtsZ-binding proteins with the GTPase domain are not required for the CTL-dependent regulation (or misregulation) of cell wall synthesis.

### The CTL of *E. coli* FtsZ has a modest effect on lateral interactions *in vitro*

While *Ec*ΔCTL by itself was unable to cause bulging and rapid lysis, the pattern of HADA localization – diffuse, asymmetric foci – suggests that *Ec*ΔCTL (*Ec*GTPase-*Ec*CTC) could still affect the organization of cell wall synthetic enzymes. We hypothesized that, similar to *Cc*ΔCTL, *Ec*ΔCTL also had aberrant assembly properties that affect downstream localization or activity of cell wall enzymes. However, due to the lack of strong protein-protein interactions through the CTC, this mutant is unable to cause bulging and lysis. To determine if CTL deletion affects FtsZ assembly properties in *E. coli* as it does in *C. crescentus*, we imaged polymers formed by *Ec*ΔCTL and compared them to *Ec*FtsZ and *Cc*ΔCTL *in vitro*. By electron microscopy, we observed that *Cc*FtsZ formed gently curved single protofilaments (Figure 3). *Ec*FtsZ, under the same conditions, predominantly formed similar gently curved single protofilaments (Figure 3). As shown previously, *Cc*ΔCTL formed straight multifilament bundles that were often longer than a micron, in addition to the structures similar to those formed by *Cc*FtsZ. *Ec*ΔCTL, on the other hand, did not form large multifilament bundles. However, *Ec*ΔCTL polymers were more dense on the grid than *Ec*FtsZ and, in addition to gently curved structures similar to *Ec*FtsZ, *Ec*ΔCTL formed straight and curved bundles that had multifilament thickness. Overall, whereas *Cc*CTL prevents the formation of long straight bundles, *Ec*CTL has a milder effect on lateral interaction. Nevertheless, the CTL appears to be important for regulating interprotofilament interactions in both *C. crescentus* and *E. coli*.

**Figure 3:**
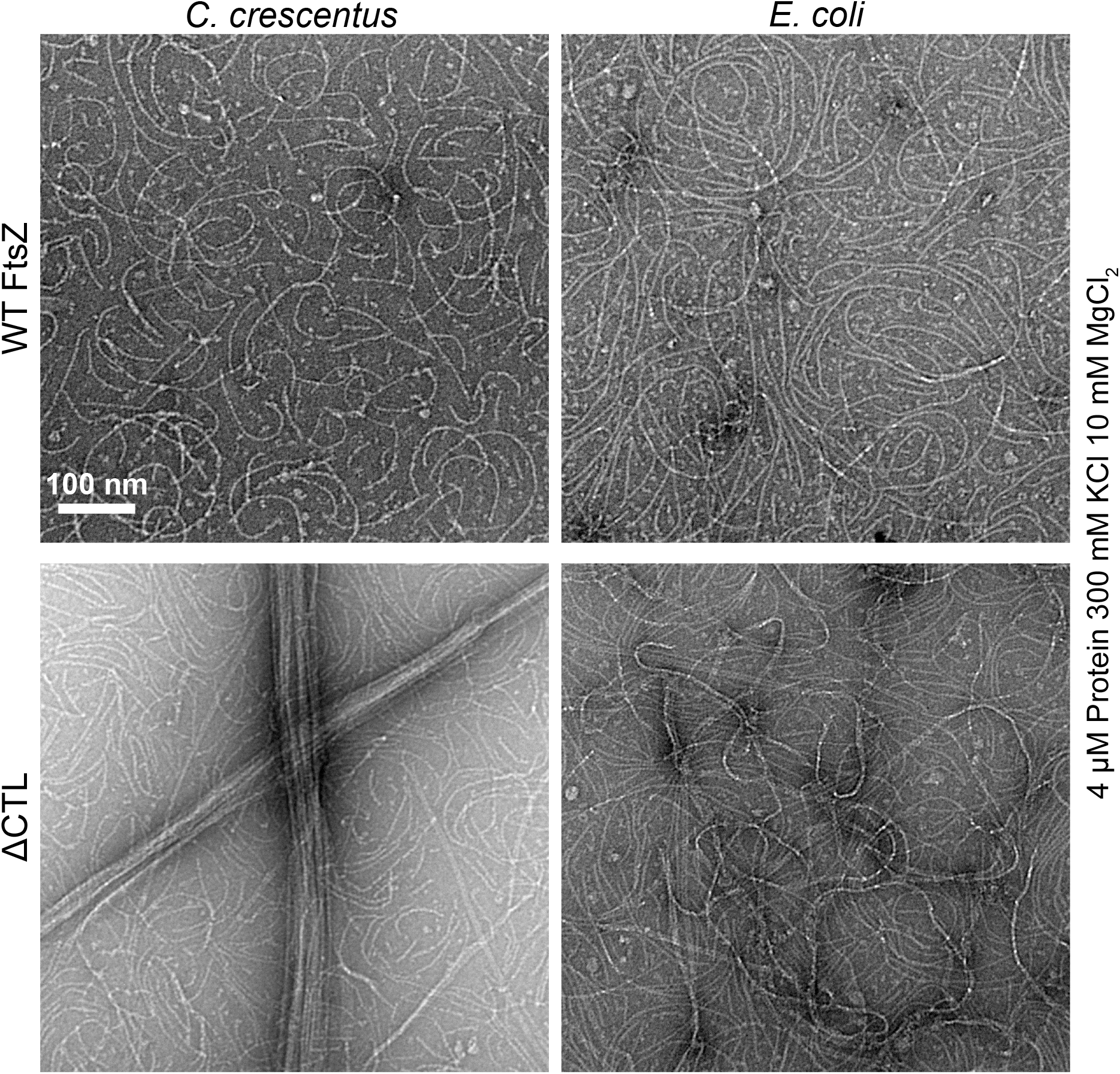
CTL regulates lateral interaction between protofilaments in *E. coli* FtsZ. Transmission electron micrographs showing polymers formed by 4 µM FtsZ or ΔCTL from *C. crescentus* or *E. coli* after incubation with GTP for 16 minutes in the presence of 300 mM KCl and 10 mM MgCl_2_. Scale bar – 100 nm.

### FtsA is required for ΔCTL-induced bulging

The ability of *Ec*GTPase-*Cc*CTC to cause bulging and lysis suggests that divisome proteins that bind to the CTC may be critical for CTL-dependent regulation of cell wall metabolism, whereas those that interact with the GTPase domain of FtsZ are likely not required. To test this, we asked if xylose-inducible expression of Δ*CTL* can cause bulging and lysis in cells deleted for non-essential members of the divisome that bind FtsZ at the GTPase domain – *zapA*-*zauP* – the CTC – *fzlC –* or at an unknown site – *ftsE*. We did not observe an effect of deleting any of these genes on ΔCTL-induced bulging or lysis (Figure 4A), indicating that CTL-dependent regulation of cell wall metabolism does not require these divisome proteins.

**Figure 4:**
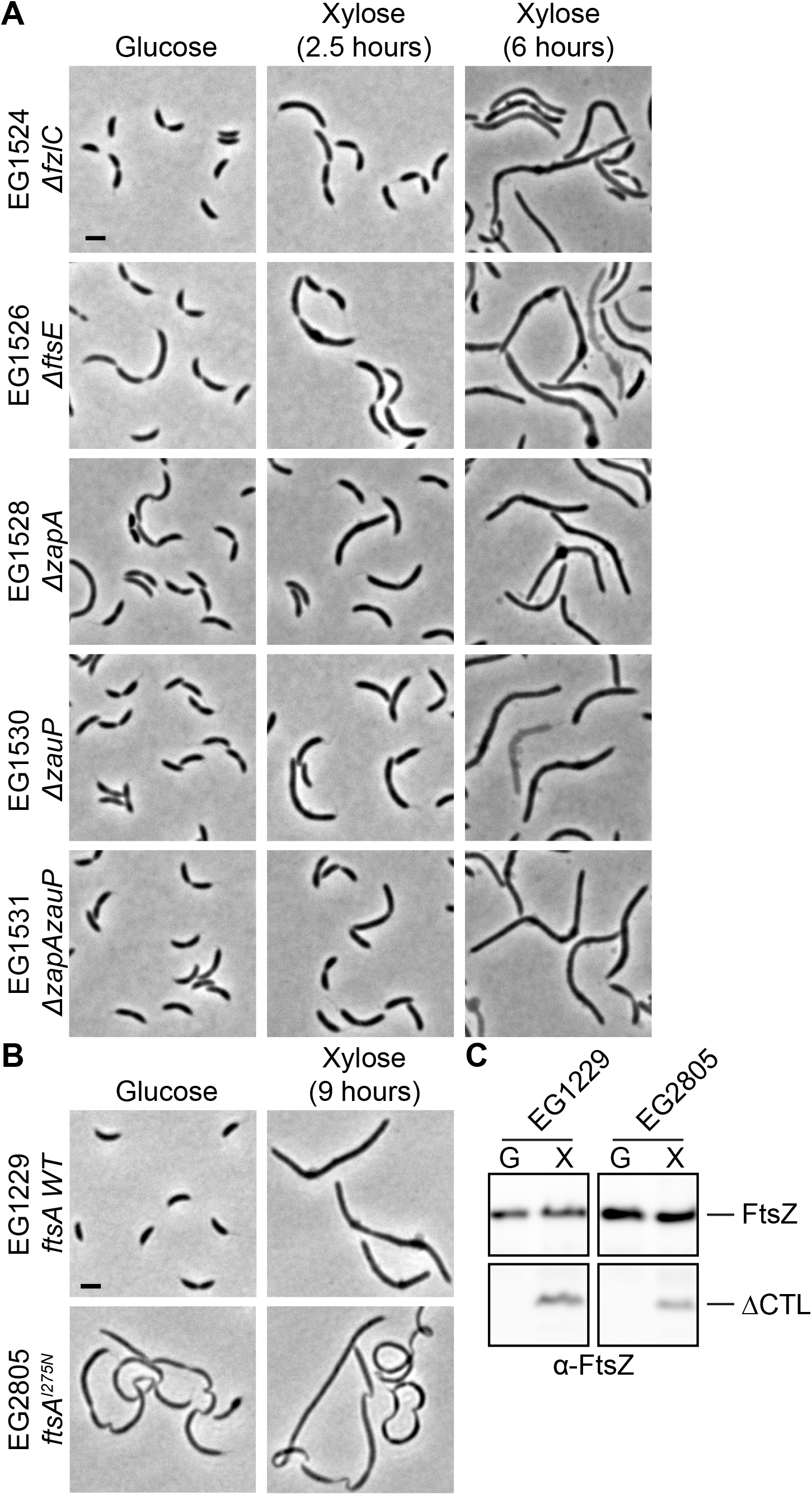
A temperature-sensitive FtsA mutant prevents ΔCTL-induced bulging. **A.** Phase contrast images showing the morphologies of cells in the absence and presence of inducer (xylose) for expression of Δ*CTL* in strains deleted for the non-essential binding partners of FtsZ. Scale bar – 2 µm. **B.** Phase contrast images showing the effects of xylose-induced expression of Δ*CTL* in cells producing a temperature sensitive allele of *ftsA* (*ftsA^I275N^*) as the only copy of *ftsA* at 30 °C. Scale bar – 2 µm. **C.** Immunoblot using anti-FtsZ antisera against lysates from cells in B. after 9 hours to confirm expression of xylose inducible Δ*CTL*. Strain key (all have xylose-inducible Δ*CTL*): Δ*fzlC* (EG1524), Δ*ftsE* (EG1526), Δ*zapA* (EG1528), Δ*zauP* (EG1530), Δ*zapAzauP* (EG1531), *ftsA^WT^* (EG1229), *ftsA^I275N^* (EG2805)

FtsA is an essential membrane-anchoring protein of FtsZ that binds to the CTC. Since FtsA is essential, we expressed Δ*CTL* in a strain (EG1776) wherein the only copy of *ftsA* is replaced by a temperature sensitive (*ftsA ts*) allele (19). At the restrictive temperature of 37°C, the *ftsA ts* allele led to filamentation in the presence or absence of ΔCTL production, but failed to exhibit characteristic bulging and lysis upon induction of Δ*CTL* (Figure S8A,D). However, this phenotype was also noted upon ΔCTL induction at the permissive temperature, prompting us to sequence the ts mutation (I275N) and clone it into a strain with an otherwise wild type background (excepting xylose-inducible Δ*CTL*, EG2805). This strain exhibited slow growth rate and filamentous morphology at 30°C and these defects are exacerbated at 37°C without induction of ΔCTL (Figure S8A, C), implying that there are additional mutations in the original temperature sensitive strain that suppress filamentation at 30°C. However, at both permissive and restrictive temperatures, EG2805 did not show any bulging upon Δ*CTL* expression (Figure 4B-C, Figure S8A,D), suggesting that wild type FtsA is required for the dominant lethal bulging effect of ΔCTL and likely plays a key role in CTL-dependent regulation of cell wall metabolism. Collectively, our data thus far indicate that the CTC is required for ΔCTL-mediated signaling through FtsA to misregulate cell wall metabolism, but that all other non-essential FtsZ-binding proteins are dispensable for this signaling.

### Overproduction of ZapA or FzlC slows the dominant lethal effects of ΔCTL

Next, we determined if the lethal effects of ΔCTL can be suppressed by increasing the concentrations of any known FtsZ-binding proteins. To this end, we overproduced known binding partners of FtsZ – FzlA, ZapA-ZauP (individually or together), MipZ, FzlC, FtsE, or both FtsE and FtsX – in cells inducibly expressing Δ*CTL*. In all the cases tested, Δ*CTL* expression led to filamentation and cell death (Figure 5, Figure S9). However, we found that overproduction of either ZapA (without ZauP) or FzlC suppressed the formation of bulges and caused initiation of constriction in cells producing ΔCTL (Figure 5A-B). Overproduction of ZapA or FzlC in the absence of ΔCTL causes a minimal increase in cell length (20–21). In cells overproducing ZapA or FzlC and producing ΔCTL, the region of constriction appeared elongated and many cells were chained. Overproduction of FtsE or FtsEX in the absence of Δ*CTL* expression causes cell elongation and ectopic pole formation (22). We observed a similar phenotype in cells overproducing FtsE or FtsEX and producing ΔCTL, with no envelope bulges, suggesting that the effects of *ftsEX* overexpression is dominant to ΔCTL-induced toxicity (Figure 5A).

**Figure 5:**
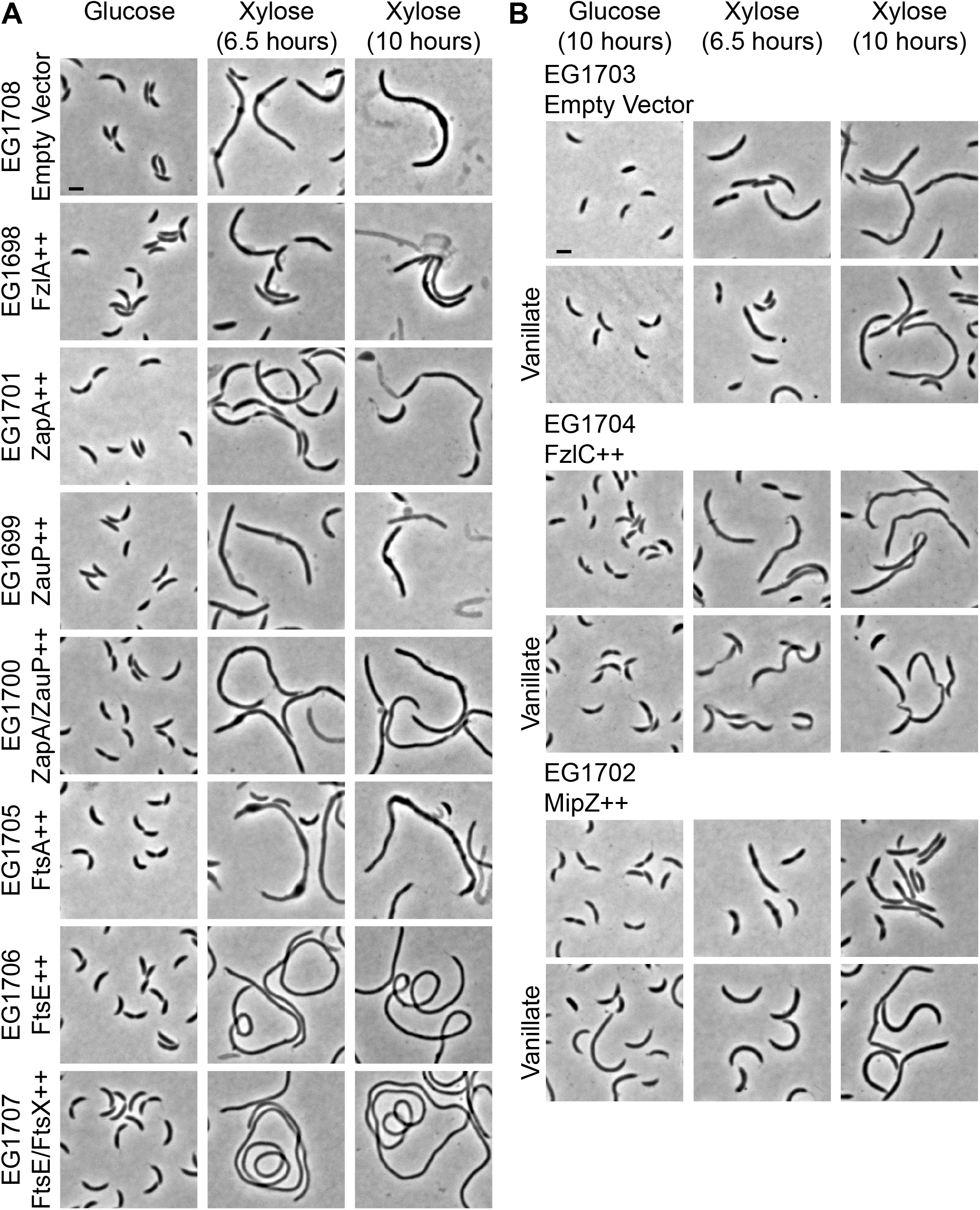
ZapA, FtsE, FzlC, or MipZ overproduction prevents formation of ΔCTL-induced bulges. **A.** Phase contrast images of cells in the absence or presence of inducer (xylose) for the expression of Δ*CTL* and overproduction of FtsZ binding proteins (or empty vector control). Scale bar – 2 µm. **B.** Phase contrast images of cells with xylose-inducible expression of Δ*CTL* and vanillate-inducible overexpression of *fzlC*, *mipZ*, or empty vector control. Scale bar – 2 µm. Strain key (all have xylose-inducible Δ*CTL*): Empty Vector control for xylose-inducible overexpression (EG1708), FzlA++ (EG1698), ZapA++ (EG1701), ZauP++ (EG1699), ZapA/ZauP++ (EG1700), FtsA++ (EG1705), FtsE++ (EG1706), FtsE/FtsX++ (EG1707), Empty Vector control for vanillate-inducible overexpression (EG1703), FzlC++ (EG1704), MipZ++ (EG1702)

MipZ is a negative regulator of FtsZ polymerization, and overproducing it inhibits assembly of Z-rings. Overexpression of MipZ in cells expressing Δ*CTL* prevented the formation of envelope bulges (Figure 5B). Since the ability of ΔCTL to form polymers is required for inducing bulging and lysis (4), overproducing MipZ could affect ΔCTL-induced bulging by preventing ΔCTL assembly. Although overproduction of ZapA, FzlC, FtsE, FtsEX, or MipZ is sufficient to suppress ΔCTL-induced bulging, none of the proteins, when overproduced, were able to support growth, as demonstrated by spot-dilution assay (Figure S9). We confirmed that the effects of overproducing these proteins on ΔCTL-induced bulging were not due to differences in expression of ΔCTL using immunoblotting (Figure S10).

In contrast to the effects of overproducing ZapA or FzlC, overproduction of FtsA caused an exacerbation of the effects of ΔCTL production – envelope bulges were larger, less symmetric, and appeared earlier in ΔCTL-producing cells overexpressing *ftsA* compared to those not overexpressing *ftsA* (Figure 5A). We conclude that of all the division proteins tested, FtsE, ZapA, FzlC, and FtsA have genetic interactions with the ΔCTL-induced bulging phenotype. Of these, only FtsA is required for ΔCTL-induced bulging.

Unlike FtsA, the membrane anchoring proteins FzlC and FtsE are dispensable for ΔCTL-induced bulging (Figure 4), but their overproduction prevents the formation of bulges in cells expressing ΔCTL (Figure 5). Since FzlC interacts with FtsZ through the CTC, it is possible that overproducing FzlC titrates away the CTC of ΔCTL from binding to FtsA and causing cell wall defects. *In vitro*, the presence of FzlC reverts the effects of CTL loss on lateral interaction between protofilaments through its interaction with FtsZ’s CTC (10). Whereas ΔCTL forms straight, extended, multi-filament bundles, ΔCTL in the presence of FzlC forms gently curved single filaments similar to WT. Thus, in addition to competitively binding the CTC and preventing ΔCTL from interacting with FtsA, FzlC overproduction might have beneficial effects on Z-ring structure and/or dynamics in the context of ΔCTL, at least in the presence of WT FtsZ. Our observations suggest that the membrane-anchoring proteins of FtsZ in *C. crescentus* have distinct roles in regulating cell wall metabolism. Considering that FtsA and FzlC both require the CTC for binding, their contributions to ΔCTL-induced bulging appear to be antagonistic. It is possible that FtsA is required for the CTL-dependent regulation of cell wall metabolism and FzlC plays a regulatory role in FtsA function or is important to signaling to other cell wall metabolic proteins downstream of FtsZ (20, 22).

### Concluding remarks

Here, we refined the molecular determinants of the dominant lethal effects of FtsZ ΔCTL on cell wall metabolism. Specifically, ΔCTL-mediated dominant lethal bulging and lysis require (i) a polymerizing GTPase domain fused to (ii) the *C. crescentus* CTC (iii) with a CTL less than 14 amino acids and (iv) WT FtsA. In addition, we observed that ΔCTL assembles into distinct polymeric structures in cells, consistent with its altered polymerization properties *in vitro*. Our study centered on genetic interactions between FtsZ-binding proteins and CTL-dependent regulation of cell wall metabolism and identified FtsA as the likely first responder to ΔCTL-induced changes in Z-ring structure. While it is unclear if FtsA forms a polymeric structure *in vivo*, self-interaction of FtsA could be relevant for cell wall misregulation downstream of ΔCTL. Interestingly, FtsA and FtsZ polymers formed *in vitro* exhibit a subunit length mismatch (23–24). FtsZ’s CTL could function as a flexible linker between FtsA and FtsZ filaments that accommodates this mismatch. In the absence of the CTL, we hypothesize that aberrant FtsZ-FtsA co-assemblies form, leading to misregulation of downstream pathways that rely on FtsZ-FtsA polymer structure and/or dynamics. *In vitro* and *in vivo* efforts to characterize FtsA self-interaction are required to further resolve the contributions of FtsA to CTL-dependent regulation of FtsZ function in cell wall metabolism and cytokinesis, in general.

## Materials and Methods

### Caulobacter crescentus *growth media and conditions*

*C. crescentus* NA1000 cells were grown at 30 °C in peptone yeast extract (PYE) media. Antibiotics concentrations used in liquid (solid) media for *C. crescentus* were as follows: gentamycin 1 (5) µg mL^−1^, kanamycin 5 (25) µg mL^−1^, spectinomycin 25 (100) µg mL^−1^, streptomycin (5 µg mL^−1^). For experiments with inducible expression of genes, inducer concentrations used were as follows: xylose – 0.3% (w/v), vanillate – 0.5 mM, glucose – 0.2% (w/v).

### Microscopy and image analysis

Cells were immobilized on 1% agarose pads and imaged using a Nikon Eclipse Ti inverted microscope through a Nikon Plan Fluor x 100 (numeric aperture 1.30) oil Ph3 objective with a Photometrics CoolSNAP HQ^2^ cooled CCD camera. Images were prepared for figure presentation in Adobe Photoshop by adjusting the fluorescence channel of each image to the same levels across samples in a given experiment (without saturating pixels or losing data) and merging on top of the corresponding phase image (shown in blue). Prior to analyzing images in Figure 1A, the background was subtracted from raw fluorescence images by finding the average value of a rectangular region of interest where no cells were present and subtracting that value from the whole image. Images were input into either Oufti (16) - for demographs and FWHM - or the MicrobeJ plugin of FIJI (15) for Z-ring fraction and intensity. Cells were then outlined with meshes using phase images and fluorescence signal was analyzed. FWHM calculations of midcell mNG signal from Oufti output were performed using a custom MATLAB script (25) which fit the normalized signal output from Oufti into an eighth term Fourier series model and determined the width of the fluorescence curve at 50% of maximum intensity. Z-rings were defined using the maxima detection function in MicrobeJ and verified manually. Mean fluorescence intensities of the Z-ring and whole cells were determined as the mean intensity within the Z-ring region of interest or whole cell, respectively. The fraction of FtsZ in the Z-ring was determined by dividing the integrated fluorescence intensity within the Z-ring by the integrated fluorescence intensity of the corresponding cell. A one-way ANOVA Kruskal-Wallis test with Dunn’s post-test was used to compare each pair of groups within each data set and determine significance.

### Spot dilution assay

Cells were grown without inducer until they reached log phase (absorption at 600 nm of 0.1 – 0.7). Then, cultures were diluted to OD_600_ of 0.05 and serially diluted up to 10^−6^ before spotting onto PYE plates containing glucose (0.2% w/v), xylose (0.3% w/v), and/or vanillate (0.5mM) as indicated (along with antibiotics corresponding to the resistance of each strain). The plates were then incubated at 30 °C until the appearance of colonies at the lowest dilution in the control strain in the glucose plates (48 hours).

### Growth rate measurement

Cells were grown until they reached log phase. They were then diluted to OD_600_ of 0.05 and inducer (or glucose control) was added at the beginning of growth measurements. OD_600_ values of three technical replicates for each culture was measured every 30 minutes for 24 hours in 96-well plates using Tecan Infinite 200 Pro plate reader with intermittent shaking and incubation at 30 °C.

### Immunoblotting

Immunoblotting for FtsZ and ΔCTL were performed using standard procedures. For anti-FtsZ blots, *Cc*FtsZ antiserum was used at 1:20,000 dilution (4) to determine levels of WT FtsZ, ΔCTL and other variants of FtsZ in lysates collected at the specified time points. Additionally, *Ec*FtsZ antiserum (a gift from Harold Erickson) was used at 1:1000 to recognize variants *E. coli* FtsZ or variants containing parts of *E. coli* FtsZ. SpmX antiserum was used as a control for loading concentration at 1:50,000 dilution (26). For anti-mNeonGreen blots, anti-mNeonGreen antibody (ChromoTek) was used at 1:1,000 dilution to determine levels of indicated mNG-FtsZ variants at specified time points. anti-Huß antibody was used as a loading control at 1:50,000 dilution (27). anti-rabbit or anti-mouse secondary antibodies conjugated to horseradish peroxidase were used at 1:10,00 dilution (PerkinElmer). Immunoblots were developed using PerkinElmer Western Lightning Plus-ECL and imaged with a GE Healthcare Amersham Imager 600. Quantification of immunoblots in Figure 1C and Figure S2E was completed in Image Lab 6.0 (Bio Rad) by manually finding bands, detecting the total volume within the region, and subtracting the background volume.

### HADA labeling

To image cell wall metabolism patterning, cells were incubated with 0.82 mM HADA for 5 minutes with shaking at 30 °C. Following incubation, cells were removed, washed twice in PBS, and resuspended in PBS before imaging. Alternately, when cells were not imaged immediately, they were fixed by resuspension in 100% ethanol and incubated on ice (at 4°C) until imaging and were pelleted and resuspended in PBS before imaging.

### Protein purification

FtsZ and ΔCTL from *C. crescentus* and *E. coli* were purified using the protocol described for *Cc*FtsZ in Sundararajan and Goley, 2017 (10). Briefly, FtsZ or ΔCTL expression was induced from pET21 vectors in *E. coli* Rosetta(DE3)pLysS using 0.5 mM IPTG at 37 °C for 3 hours after the uninduced cultures reached an OD_600_ of 1.0. Cells were then pelleted and resuspended in lysis buffer (50 mM Tris-HCl pH 8.0, 50 mM KCl, 1 mM EDTA, 10% glycerol, DNase I, 1 mM β-mercaptoethanol, 2 mM PMSF, 1 cOmplete mini, EDTA-free Protease inhibitor tablet (Roche)). Resuspended cell pellets were lysed by incubation with 1 mg/mL lysozyme for 1 hour followed by sonication. FtsZ or ΔCTL were then purified from the lysate using anion exchange chromatography column (HiTrap Q HP 5 ml, GE Life Sciences), followed by ammonium sulfate precipitation (at 20% - 30% ammonium sulfate saturation). The ammonium sulfate pellet was resuspended in FtsZ storage buffer (50 mM HEPES KOH pH 7.2, 50 mM KCl, 0.1 mM EDTA, 1 mM β-mercaptoethanol, 10% glycerol) and was subjected to size-exclusion chromatography (Superdex 200 10/300 GL, GE Life Sciences) to further purify the protein and was snap frozen in liquid nitrogen and stored in FtsZ storage buffer at −80 °C.

### Transmission electron microscopy

TEM of FtsZ or ΔCTL polymers were performed according to the protocol described in Sundararajan and Goley, 2017. 4 µM protein was incubated with 2 mM GTP in buffer containing 50 mM HEPES-KOH pH 7.2, 300 mM KCl, 10 mM MgCl_2_, 0.1 mM EDTA for 16 minutes prior to spotting on glow-discharged carbon coated grids. The grids were then blotted and incubated for 2 minutes with 0.75% Uranyl Formate solution twice and air dried. The stained grids were imaged using a Philips/FEI BioTwin CM120 TEM (operated at 80 kV) equipped with an AMT XR80 8 megapixel CCD camera (AMT Imaging, USA).

## Acknowledgements

We thank members of the Goley lab for helpful discussions throughout this work and feedback on the manuscript. We are grateful to Patrick Viollier, Martin Thanbichler, Lucy Shapiro, and Harold Erickson for antibodies and strains and to Caren Freel Meyers and Amer Al-khouja for synthesis of HADA. This work funded in part by the National Institutes of Health, National Institute of General Medical Sciences through R01GM108640 (EDG) and T32GM007445 (training grant support of KS and JMB).

## Author Contributions

KS and EDG conceived the study. All authors designed and carried out experiments, analyzed data, and contributed to writing and editing the manuscript.

