## Supplemental Material for "Determinants of FtsZ C-terminal linker-dependent regulation of cell wall metabolism in *Caulobacter crescentus*"

**Supplemental figure legends:**

**Figure S1: Domain architecture of FtsZ CTL variants**

Domain structures of FtsZ, ΔCTL, L14, and *Hn*CTL variants used in this study. Numbers indicated correspond to *Cc*FtsZ amino acid sequence.

**Figure S2: ΔCTL assembles into large asymmetric superstructures at sites of cell wall bulging in cells depleted of WT FtsZ.**

**A.-B.** Phase contrast, epifluorescence, and merged images of cells induced with xylose to drive expression of *mNG-FtsZ, mNG-ΔCTL, mNG-L14,* or *mNG*-Hn*CTL* from $P_{xylX}$ promoter for 2.5 hours (**A**) or 5 hours (**B**) while simultaneously depleting WT FtsZ. Scale bar – 2 μm. Strain key: mNG-FtsZ (EG2095), mNG-ΔCTL (EG2096), mNG-L14 (EG2097), mNG-*Hn*CTL (EG2098)

**Figure S3: Demographs show differences in FtsZ distribution among CTL variants in FtsZ depletion and FtsZ WT backgrounds.**

**A.-B.** Demographs showing mNG intensity as a function of cell length (blue = least intense pixel in each cell, red = most intense pixel in each cell) of a population of cells (length ≤ 5μm) represented in Figure 1A (**A**) and Figure S5A (**B**). Strain key: mNG-FtsZ (EG2095), mNG-ΔCTL (EG2096), mNG-L14 (EG2097), mNG-*Hn*CTL (EG2098), mNG-FtsZ/FtsZ (EG2102), mNG-ΔCTL/FtsZ (EG2103), mNG-L14/FtsZ (EG2104), mNG-*Hn*CTL/FtsZ (EG2105)

**Figure S4: mNG-ΔCTL and mNG-L14 FtsZ variants are present at elevated levels in cells depleted of WT FtsZ.**

**A.-B.** Immunoblots using anti-mNeonGreen antibody against lysates from cells depleted of FtsZ and uninduced (glucose and vanillate) or induced (xylose) for *mNG-FtsZ, mNG-ΔCTL, mNG-*

*L14,* or *mNG*-Hn*CTL* for 1 hour (**A**) or 4.5 hours (**B**). Overexposed blots are also shown below each blot to validate lack of signal in uninduced controls. Anti-Huß antibody was used as concentration control for loading and quantification. **C.** Quantification of immunoblots from A. and B. showing relative abundance of mNG-FtsZ variants after 1 hour (black) or 4.5 hours (grey). Bars represent standard deviation. * - $P \leq 0.05$; ** - $P \leq 0.01$; *** - $P \leq 0.001$. Strain key: mNG-FtsZ (EG2095), mNG-ΔCTL (EG2096), mNG-L14 (EG2097), mNG-*Hn*CTL (EG2098)

**Figure S5: The CTL impacts Z-ring organization in the presence of WT FtsZ.**

**A.-B.** Phase contrast, epifluorescence, and merged images of cells induced with xylose to drive expression of *mNG-FtsZ, mNG-ΔCTL, mNG-L14,* or *mNG*-Hn*CTL* from the $P_{xylX}$ promoter for 1 hour (**A**) or 5 hours (**B**) in strains producing WT FtsZ. **C.-F.** Quantification of epifluorescence images of cells 3 to 5 µm long indicating the full-width at half max (FWHM) values of Z-ring intensity (**C),** fraction of mNG-FtsZ or variants in the Z-ring (**D**), and mean epifluorescence intensity of the Z-ring (**E**) or the entire cells (**F**) in a WT FtsZ background. Bars represent standard deviation. * - $P \leq 0.05$; ** - $P \leq 0.01$; *** - $P \leq 0.001$. Strain key: mNG-FtsZ/FtsZ (EG2102), mNG-ΔCTL/FtsZ (EG2103), mNG-L14/FtsZ (EG2104), mNG-*Hn*CTL/FtsZ (EG2105)

**Figure S6: mNG-ΔCTL FtsZ is present at elevated levels in the presence of WT FtsZ.**

**A.-B.** Immunoblots using anti-mNeonGreen antibody against lysates from cells uninduced (glucose) or induced (xylose) for *mNG-FtsZ, mNG-ΔCTL, mNG-L14,* or *mNG*-Hn*CTL* for 1 hour (**A**) or 4.5 hours (**B**). Overexposed blots are also shown below each blot to validate lack of signal in uninduced controls. Anti-Huß antibody was used as concentration control for loading and quantification. **C.** Quantification of immunoblots from A. and B. showing relative abundance of mNG-FtsZ variants after 1 hour (black) or 4.5 hours (grey). Bars represent standard deviation. * - $P \leq 0.05$. Strain key: mNG-FtsZ/FtsZ (EG2102), mNG-ΔCTL/FtsZ (EG2103), mNG-L14/FtsZ (EG2104), mNG-*Hn*CTL/FtsZ (EG2105)

**Figure S7: FtsZ chimeras are produced to similar steady state levels.**

**A.-B.** Immunoblots using anti-*Cc*FtsZ (**A**) and anti-*Ec*FtsZ (**B**) antibodies showing levels of chimeric FtsZ variants shown in Figure 2A. VG – vanillate+glucose control with only WT Cc*FtsZ* expression. X – xylose driven expression of FtsZ chimeras for 5 hours. Anti-SpmX antibody was used as concentration control for loading. Asterisk indicates a non-specific band. Strain key: *Cc*FtsZ (EG951), *Cc*ΔCTL (EG852), *Cc*L14 (EG968), *Ec*FtsZ (EG1521), *Ec*ΔCTL (EG1520), *Ec*GTPase-*Cc*CTC (EG1519), *Ec*GTPase-*Ec*CTL-*Cc*CTC (EG1522), *Ec*GTPase-*Cc*CTL-*Cc*CTC (EG1517), *Ec*GTPase-*Cc*L14-*Cc*CTC (EG1516)

**Figure S8: Temperature sensitive FtsA suppresses formation of ΔCTL-induced bulges.**

**A.** Phase contrast images of cells with WT *ftsA* or a temperature sensitive allele of *ftsA* in its original mutagenized background (*ftsA ts*) or in an otherwise wild type background (*ftsA ts**) uninduced (glucose) or induced (xylose) for Δ*CTL* at 30˚C or 37˚C for 9 hours. **B.** Immunoblot using anti-FtsZ antibody on lysates from strains from A. uninduced (G) or induced (X) for Δ*CTL* expression at 30˚C or 37˚C for 9 hours. **C.** Growth characteristics of the strains from A. uninduced (G) or induced (X) for Δ*CTL* expression and grown for 24 hours at 30˚C. **D.** Spot dilutions of strains from A. Cells in log phase were diluted to an $OD_{600}$ of 0.05, serially diluted, and spotted onto PYE agar plates with indicated inducer. Plates were incubated at 30˚C for 48 hours before imaging. Strain key (all have xylose-inducible Δ*CTL*): $ftsA^{WT}$ (EG1229), *ftsA ts* (EG1776), $ftsA^{I275N}$ (EG2805)

**Figure S9: Overexpression of FtsZ binding partners does not suppress ΔCTL-induced growth defects.**

**A.-B.** Spot dilutions of strains in Figure 4 and Figure 5 showing growth of cells uninduced (glucose) or induced (xylose) for Δ*CTL* expression. Cells in log phase were diluted to an $OD_{600}$ of 0.05, serially diluted, and spotted onto PYE agar plates with indicated inducer (glucose, xylose, and/or vanillate). Plates were incubated at 30˚C for 48 hours before imaging. Strain key (all have xylose-inducible Δ*CTL*): Empty Vector control for xylose-inducible overexpression (EG1708), FzlA++ (EG1698), ZapA++ (EG1701), ZauP++ (EG1699), ZapA/ZauP++ (EG1700), FtsA++ (EG1705), FtsE++ (EG1706), FtsE/FtsX++ (EG1707), Empty Vector control for vanillate-inducible overexpression (EG1703), FzlC++ (EG1704), MipZ++ (EG1702)

**Figure S10: Levels of FtsZ and ΔCTL are not impacted by overexpression of binding partners.**

Immunoblots using anti-FtsZ antibody showing protein levels of ΔCTL and WT FtsZ corresponding to the experiments in Figure 5 at 6.5 hours of incubation with inducers (glucose or X, vanillate or V, xylose or X). Strain key (all have xylose-inducible Δ*CTL*): Empty Vector control for xylose-inducible overexpression (EG1708), FzlA++ (EG1698), ZapA++ (EG1701), ZauP++ (EG1699), ZapA/ZauP++ (EG1700), FtsA++ (EG1705), FtsE++ (EG1706), FtsE/FtsX++ (EG1707), Empty Vector control for vanillate-inducible overexpression (EG1703), FzlC++ (EG1704), MipZ++ (EG1702)

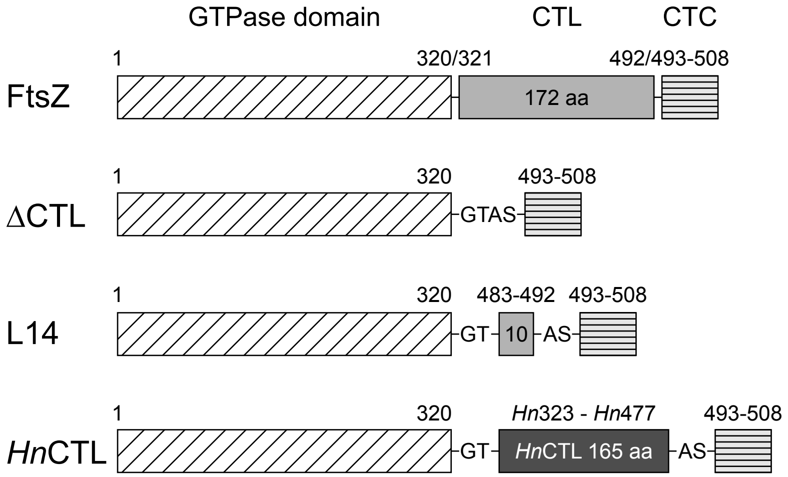
GTPase domain
CTL
CTC
1
320/321
492/493-508
FtsZ
172 aa
1
320
493-508
ΔCTL
GTAS
1
320
483-492
493-508
L14
GT
10
AS
1
320
*Hn*323 - *Hn*477
493-508
*Hn*CTL
GT
*Hn*CTL 165 aa
AS

Figure S1

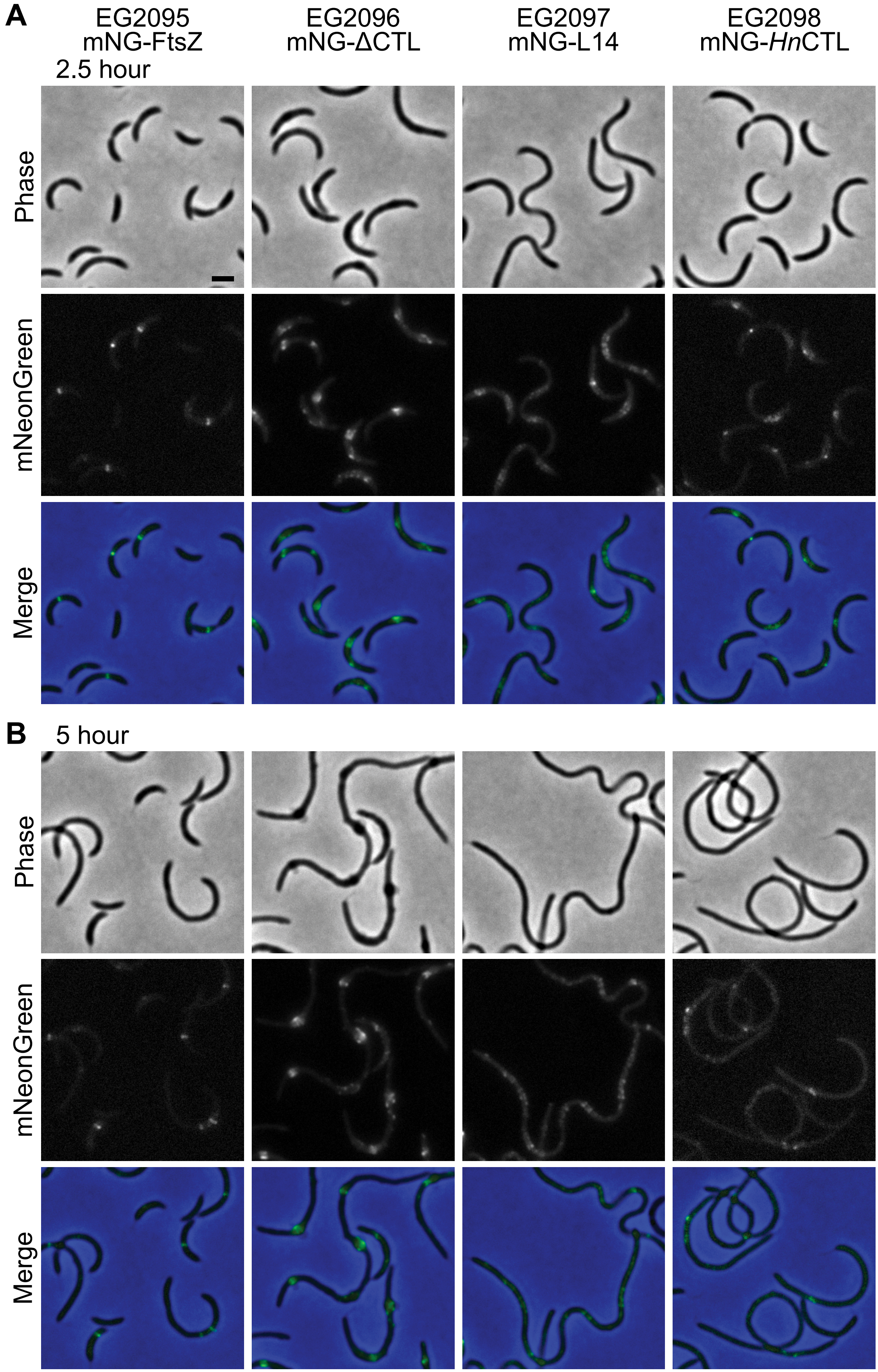
A
EG2095
mNG-FtsZ
EG2096
mNG-ΔCTL
EG2097
mNG-L14
EG2098
mNG-*Hn*CTL
2.5 hour
Phase
mNeonGreen
Merge
B
5 hour
Phase
mNeonGreen
Merge

Figure S2

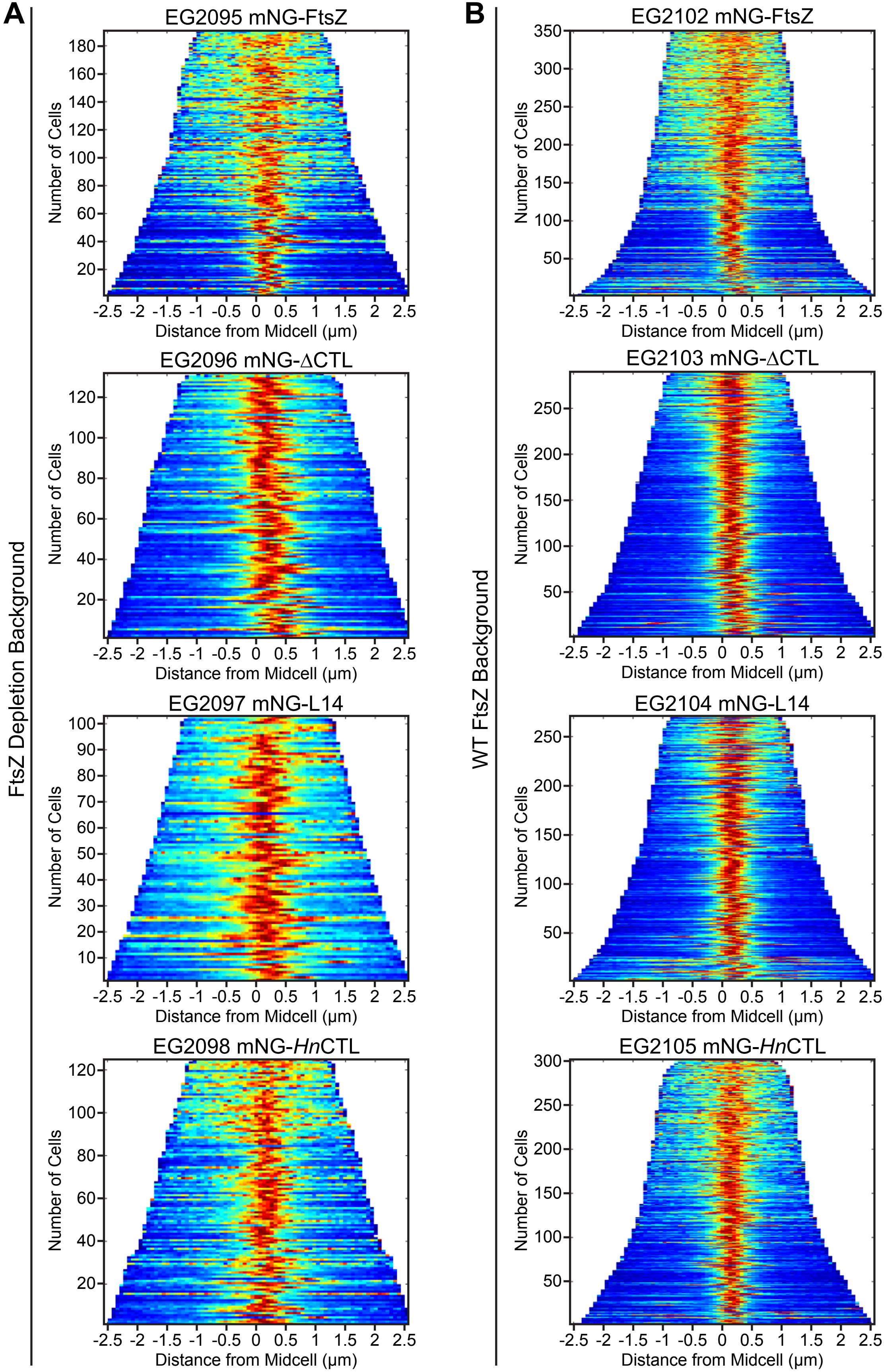

A
B
FtsZ Depletion Background
WT FtsZ Background
EG2095 mNG-FtsZ
EG2096 mNG-ΔCTL
EG2097 mNG-L14
EG2098 mNG-*Hn*CTL
EG2102 mNG-FtsZ
EG2103 mNG-ΔCTL
EG2104 mNG-L14
EG2105 mNG-*Hn*CTL
Number of Cells
Distance from Midcell (μm)


Figure S3

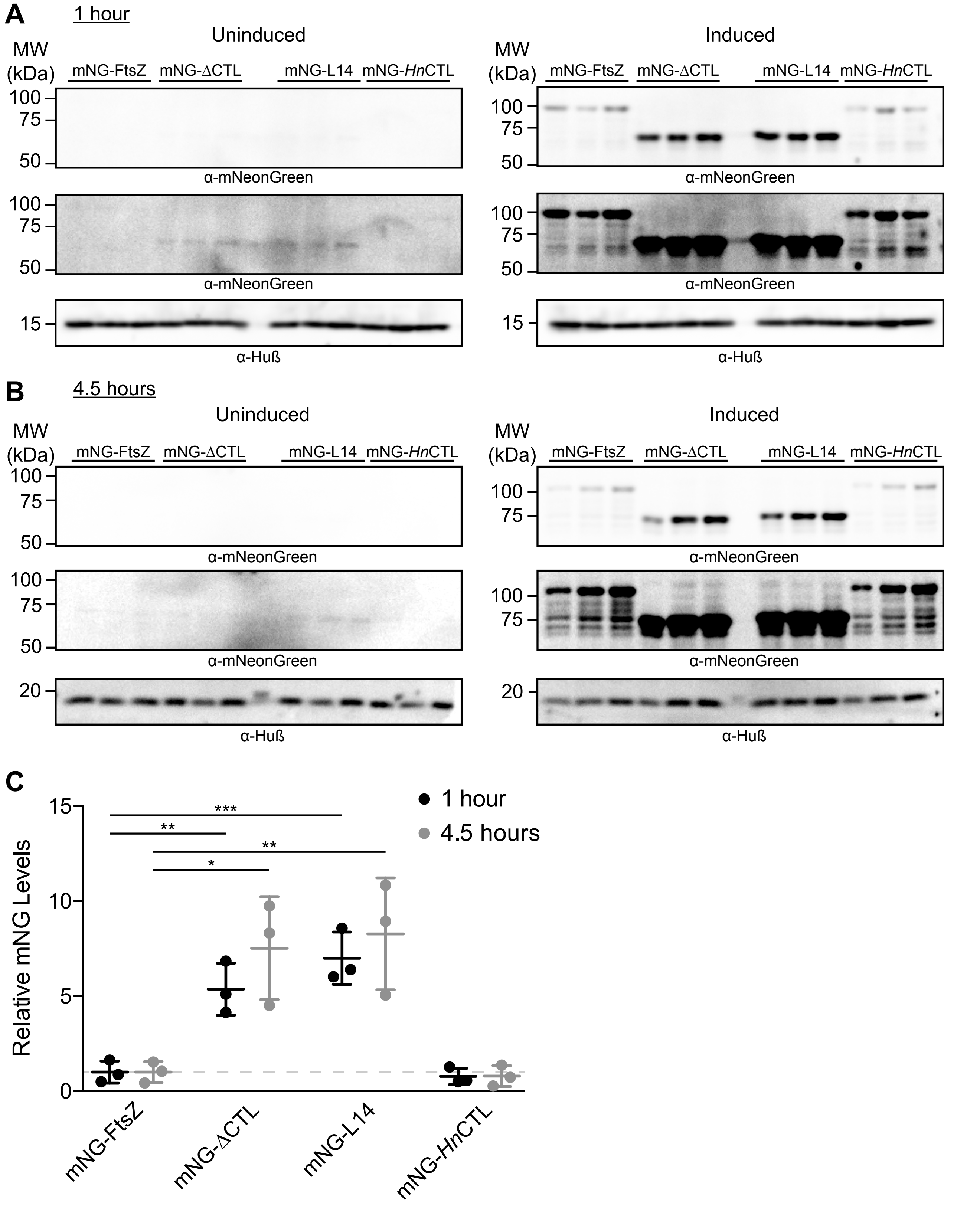
A
1 hour
Uninduced
Induced
MW (kDa)
mNG-FtsZ
mNG-ΔCTL
mNG-L14
mNG-*Hn*CTL
100
75
50
15
20
α-mNeonGreen
α-Huß
B
4.5 hours
C
Relative mNG Levels
15
10
5
0
***
**
*
1 hour
4.5 hours

Figure S4

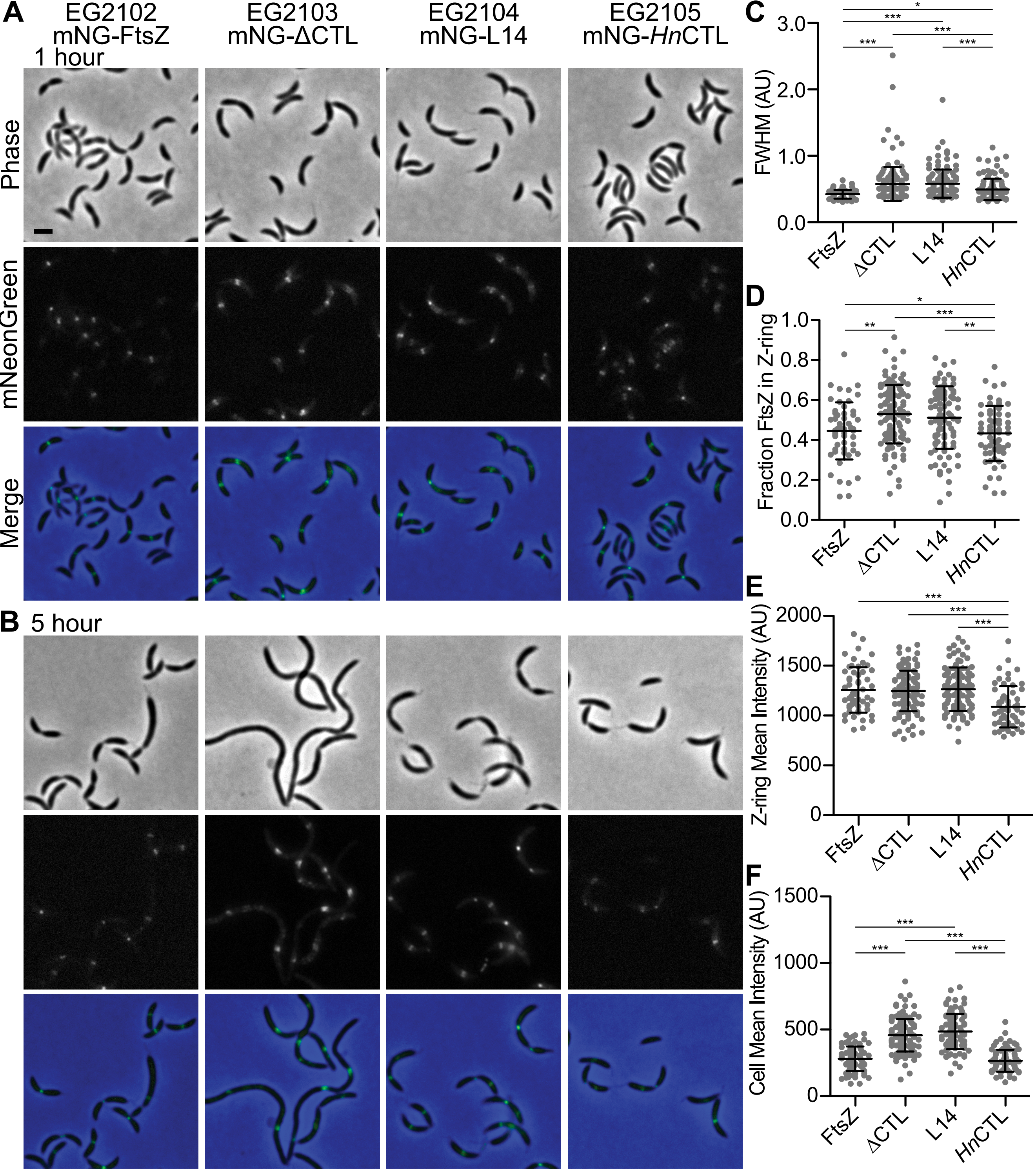
A
EG2102 mNG-FtsZ
EG2103 mNG-ΔCTL
EG2104 mNG-L14
EG2105 mNG-HnCTL
1 hour
Phase
mNeonGreen
Merge
B 5 hour
C
FWHM (AU)
D
Fraction FtsZ in Z-ring
E
Z-ring Mean Intensity (AU)
F
Cell Mean Intensity (AU)
FtsZ
ΔCTL
L14
HnCTL

Figure S5

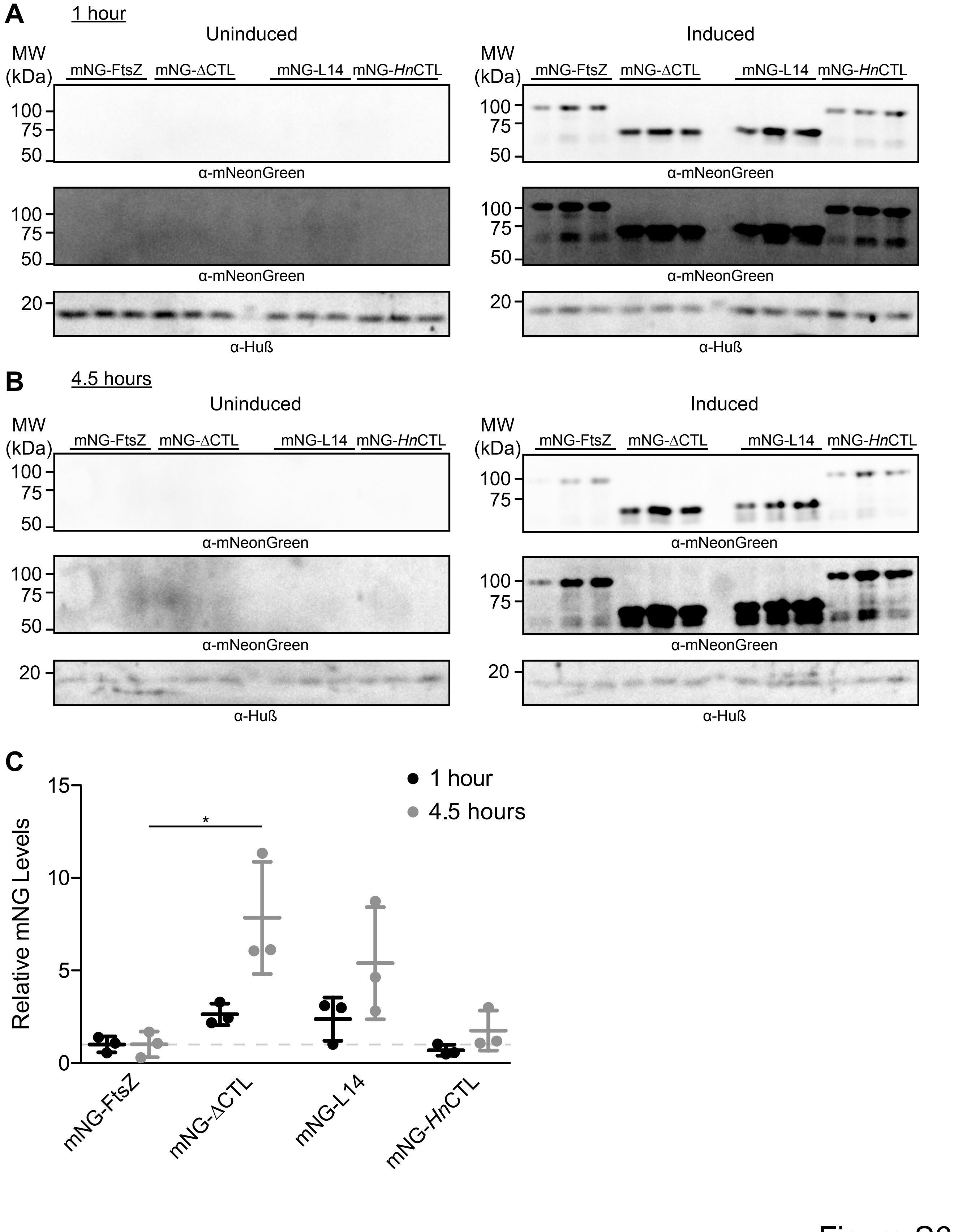
A
1 hour
Uninduced
Induced
MW (kDa)
mNG-FtsZ
mNG-ΔCTL
mNG-L14
mNG-HnCTL
100
75
50
20
α-mNeonGreen
α-Huß
B
4.5 hours
C
1 hour
4.5 hours
Relative mNG Levels
15
10
5
0
*

Figure S6

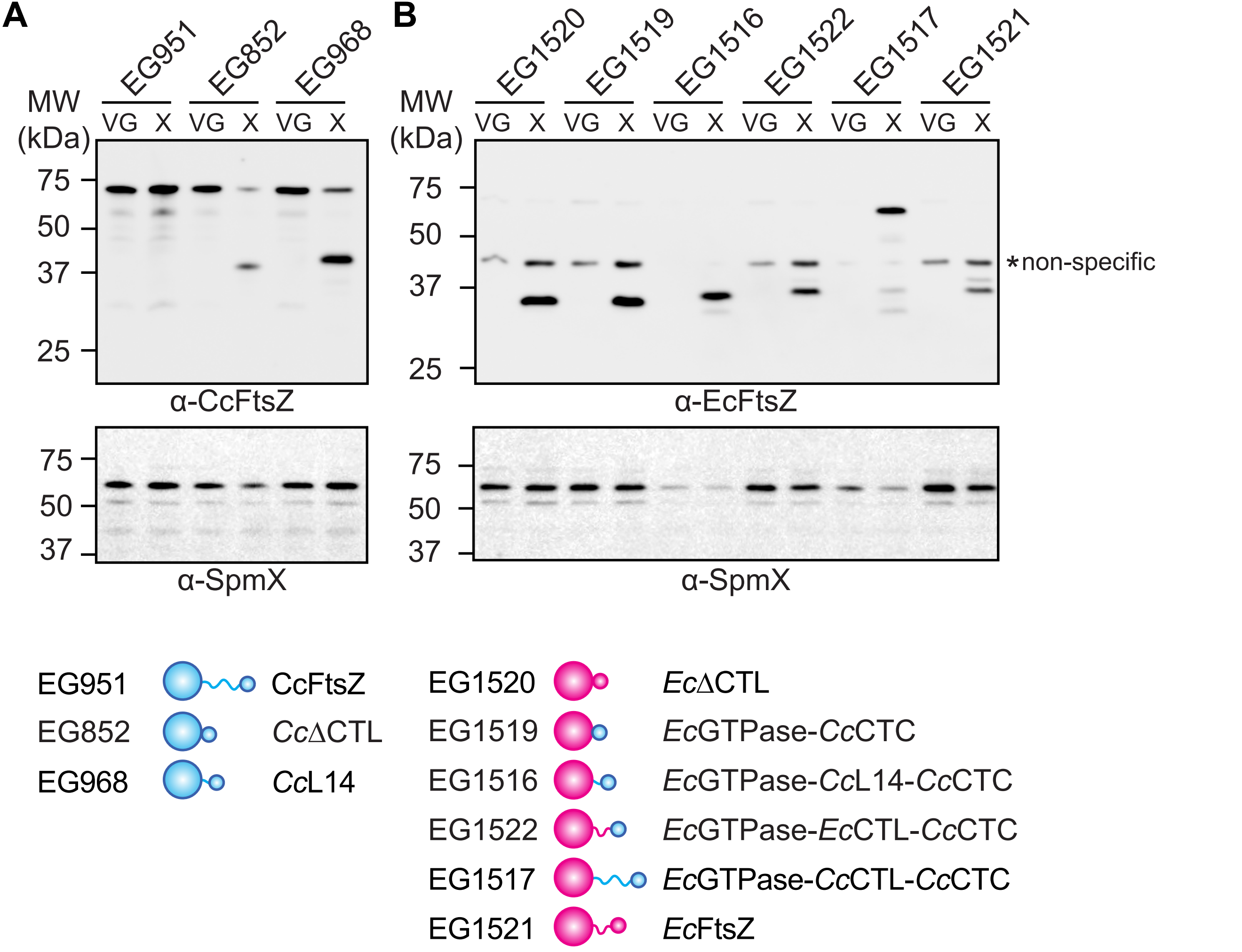
A
B
EG951
EG852
EG968
EG1520
EG1519
EG1516
EG1522
EG1517
EG1521
MW (kDa)
VG X VG X VG X
VG X VG X VG X VG X VG X VG X
75
50
37
25
α-CcFtsZ
α-EcFtsZ
*non-specific
α-SpmX
EG951 CcFtsZ
EG852 CcΔCTL
EG968 CcL14
EG1520 EcΔCTL
EG1519 EcGTPase-CcCTC
EG1516 EcGTPase-CcL14-CcCTC
EG1522 EcGTPase-EcCTL-CcCTC
EG1517 EcGTPase-CcCTL-CcCTC
EG1521 EcFtsZ

Figure S7

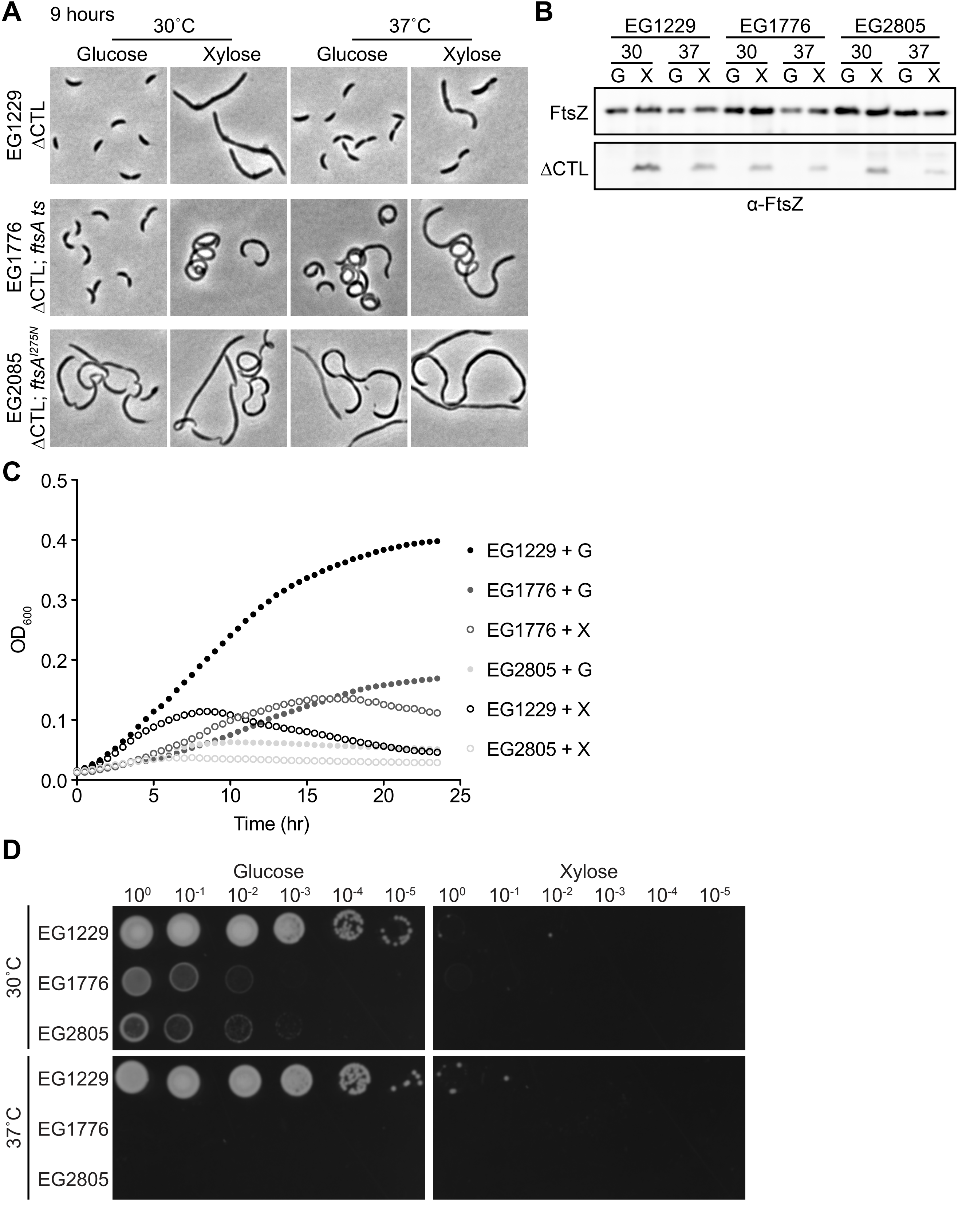
A
9 hours
30˚C
37˚C
Glucose
Xylose
Glucose
Xylose
EG1229 ΔCTL
EG1776 ΔCTL; ftsA ts
EG2085 ΔCTL; ftsA I275N
B
EG1229
EG1776
EG2805
30
37
G
X
FtsZ
ΔCTL
α-FtsZ
C
OD600
Time (hr)
EG1229 + G
EG1776 + G
EG1776 + X
EG2805 + G
EG1229 + X
EG2805 + X
D
Glucose
Xylose
30˚C
37˚C
EG1229
EG1776
EG2805

Figure S8

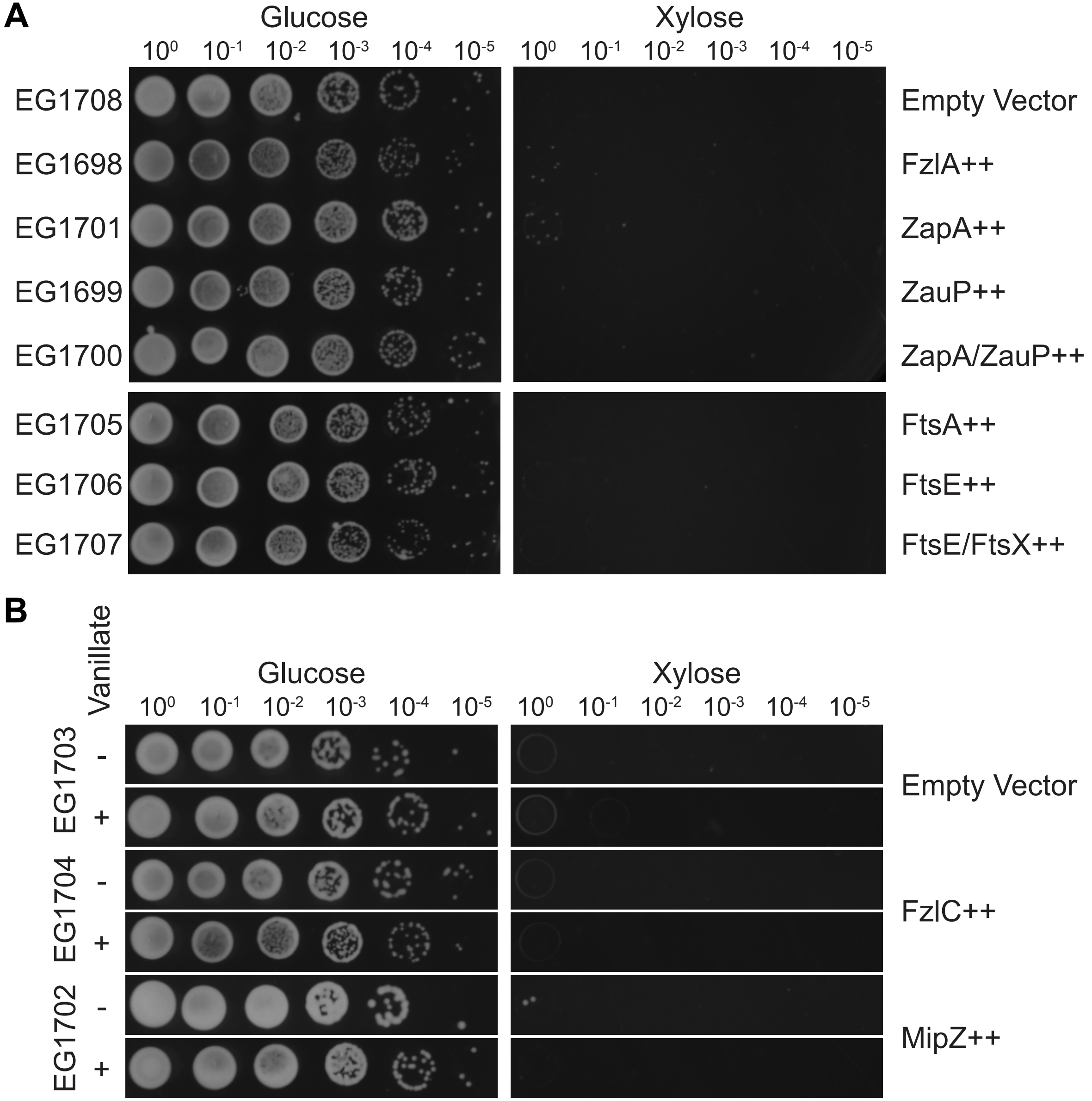
A
Glucose
Xylose
$10^0$ $10^{-1}$ $10^{-2}$ $10^{-3}$ $10^{-4}$ $10^{-5}$
EG1708
EG1698
EG1701
EG1699
EG1700
EG1705
EG1706
EG1707
Empty Vector
FzlA++
ZapA++
ZauP++
ZapA/ZauP++
FtsA++
FtsE++
FtsE/FtsX++
B
Vanillate
EG1703
EG1704
EG1702
-
+
Empty Vector
FzlC++
MipZ++

Figure S9

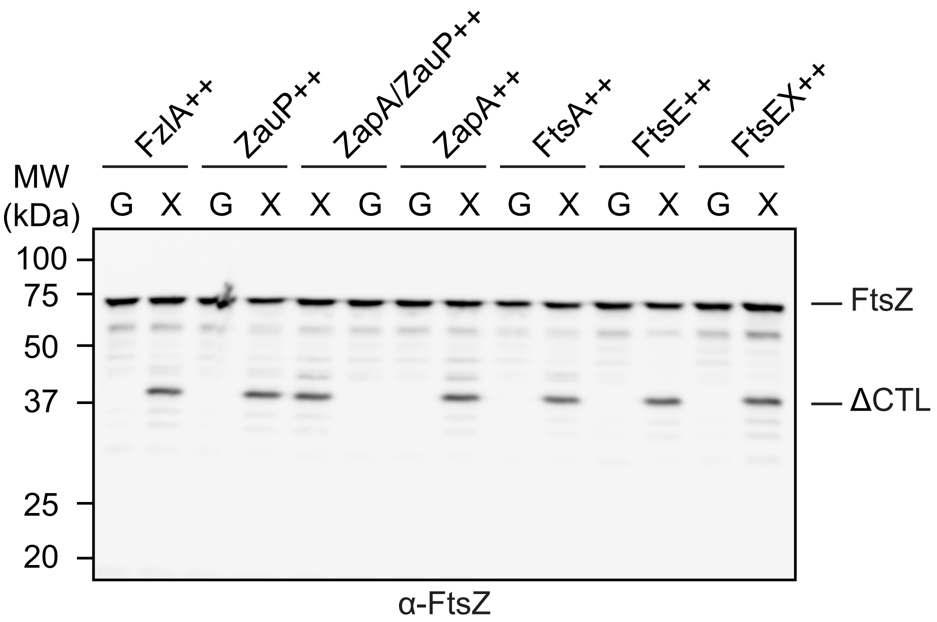
FzlA++
ZauP++
ZapA/ZauP++
ZapA++
FtsA++
FtsE++
FtsEX++
MW (kDa)
G X G X X G G X G X G X G X
100
75
50
37
25
20
— FtsZ
— ΔCTL
α-FtsZ

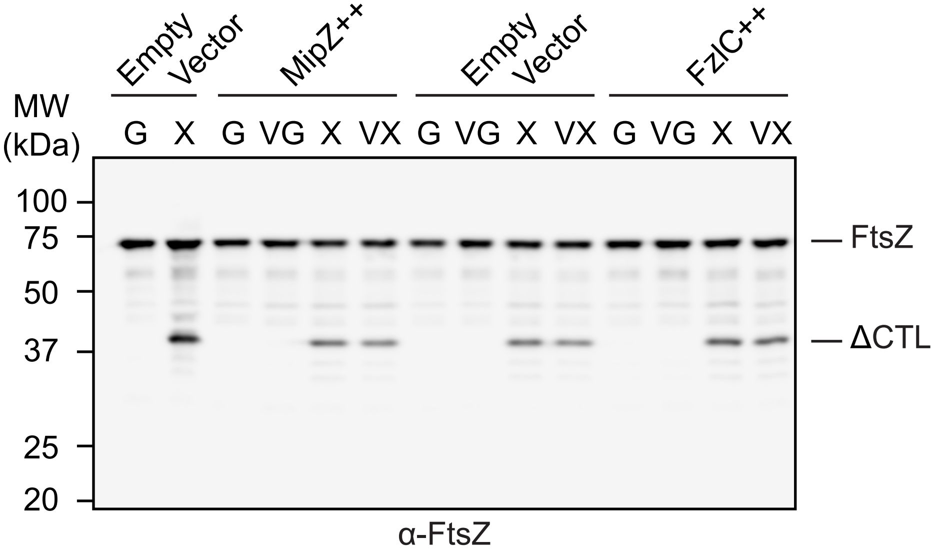
Empty Vector
MipZ++
Empty Vector
FzlC++
MW (kDa)
G X G VG X VX G VG X VX G VG X VX
100
75
50
37
25
20
— FtsZ
— ΔCTL
α-FtsZ

Figure S10
